# MYC and HSF1 Cooperate to Drive PLK1 inhibitor Sensitivity in High Grade Serous Ovarian Cancer

**DOI:** 10.1101/2024.06.11.598486

**Authors:** Imade Williams, Haddie DeHart, Matthew O’Malley, Bobby Walker, Vrushabh Ulhaskumar, Haimanti Ray, Joe R. Delaney, Kenneth P. Nephew, Richard L. Carpenter

**Author notes:** **Additional Information:** <u>Financial Support:</u> National Cancer Institute K22CA207575 (RLC); Van Andel Institute through the Van Andel Institute – Stand Up To Cancer Epigenetics Dream Team. Stand Up To Cancer is a division of the Entertainment Industry Foundation, administered by AACR (RLC, KPN); Mentored Investigator Grant from Ovarian Cancer Research Alliance (IW). <u>Corresponding Author Contact Information:</u> Address: 308 Biology Building, 1001 E. 3^rd^ St., Bloomington, IN 47405.

## Abstract

Ovarian cancer is a deadly female cancer with high rates of recurrence. The primary treatment strategy for patients is platinum-based therapy regimens that almost universally develop resistance. Consequently, new therapeutic avenues are needed to overcome the plateau that current therapies have on patient outcomes. We describe a gene amplification involving both HSF1 and MYC, wherein these two genes on chromosome 8q are co-amplified in over 7% of human tumors that is enriched to over 30% of patients with ovarian cancer. We further found that HSF1 and MYC transcriptional activity is correlated in human tumors and ovarian cancer cell lines, suggesting they may cooperate in ovarian cancer cells. CUT&RUN for HSF1 and MYC in co-amplified ovarian cancer cells revealed that HSF1 and MYC have overlapping binding at a substantial number of locations throughout the genome where their binding peaks are near identical. Consistent with these data, a protein-protein interaction between HSF1 and MYC was detected in ovarian cancer cells, implying these two transcription factors have a molecular cooperation. Further supporting their cooperation, growth of HSF1-MYC co-amplified ovarian cancer cells were found to be dependent on both HSF1 and MYC. In an attempt to identify a therapeutic target that could take advantage of this dependency on both HSF1 and MYC, PLK1 was identified as being correlated with HSF1 and MYC in primary human tumor specimens, consistent with a previously established effect of PLK1 on HSF1 and MYC protein levels. Targeting PLK1 with the compound volasertib (BI-6727) revealed a greater than 200-fold increased potency of volasertib in HSF1-MYC co-amplified ovarian cancer cells compared to ovarian cancer cells wild-type HSF1 and MYC copy number, which extended to several growth assays, including spheroid growth. Volasertib, and other PLK1 inhibitors, have not shown great success in clinical trials and this study suggests that targeting PLK1 may be viable in a precision medicine approach using HSF1-MYC co-amplification as a biomarker for response.

## Introduction

Ovarian cancer is the 5^th^ leading cause of cancer-related death in women in the US and has a dismal prognosis (1) with high-grade serous ovarian cancer (HGSOC) subtype accounting for 70%-80% of ovarian cancer deaths (2–7). High grade serous is a chemotherapy responsive tumor with high initial response rates to standard therapy consisting of platinum (Pt) and paclitaxel. However, most women eventually develop recurrence and chemotherapy-resistant disease. Recurrent ovarian cancer is essentially incurable (7). There is an urgent need for novel and improved therapies for HGSOC (2). In particular, precision medicine approaches to identify subgroups of patients that have a higher likelihood of responding to any given therapy may be critical to overcome the current plateau of patient outcomes for this disease.

Heat shock factor 1 (HSF1) is a transcription factor that was originally discovered as the master regulator of the heat shock response (8–10). This physiological role of HSF1 includes the transcriptional upregulation of chaperone heat shock proteins in response to cellular stressors (11). HSF1 was identified to be more active in basal or triple-negative breast cancers (12), which have striking similarities to HGSOC, and HSF1 was recently shown to overexpressed and/or hyperactivated in HGSOC and other solid tumors (13–38). High HSF1 activity has previously been associated with poor patient outcomes in several cancer types (12,29,33,39,40).

The oncogenic transcription factor MYC has been reported to be overexpressed at the RNA level in up to 60% of ovarian tumors (41–48). In HGSOC, MYC has been shown to promote several mechanisms that induce oncogenesis and promote cancer progression, including proliferation and tumor cell metabolism (49). MYC expression is also a reported prognostic marker for response to chemotherapy in HGSOC patients (45,50–54). There was a recent report that MYC can cooperate with HSF1in non-cancer cells (55) and in hepatocellular carcinoma (56). However, whether they have any cooperation or interact in HGSOC has not been tested.

Polo-like kinase 1 (PLK1) is a serine/threonine kinase that has functions in cell cycle progression, DNA damage response, and replication stress (57–60). PLK1 has been shown to be overexpressed and associated with poor patient outcomes in several human cancers, including HGSOC (61–68). PLK1 is a therapeutic target and four compounds targeting PLK1 have been examined clinical trials, including volasertib (BI-6727) (58). Despite a clear therapeutic window for PLK1 inhibition, the efficacy of these compounds has been limited by treatment toxicities (58). However, PLK1 has been shown to directly phosphorylate both MYC and HSF1 to enhance their activity and protein stabilization (69–73). Towards a precision medicine approach that can identify patients who would benefit from PLK1 inhibition, we have identified a therapeutic vulnerability of HSF1-MYC co-amplification, which accounts for approximately one-third of ovarian cancer patients and has significance for other solid tumors, including breast cancer. We demonstrate that these two transcription factors have a functional interaction and cooperation in HGSOC cells. Furthermore, HGSOC cells with HSF1 and MYC co-amplification are more sensitive to PLK1 inhibition than cells with normal copy numbers of HSF1 and MYC.

## Methods and Materials

### TCGA Analysis

Gene amplification status in all TCGA cohorts was determined using called amplification status (≥2 copy gain) from publicly available TCGA data in cBioPortal.org. RNA-sequencing of the TCGA ovarian cancer cohort (TCGA-OV) were downloaded in RPKM for included analyses. MYC activity was assessed using a published gene signature (74) along with HSF1 activity (12).

### Cell Culture and Reagents

Unless otherwise indicated, all cell lines were purchased from ATCC and cultured in ATCC-recommended culture media at 37°C with 5% CO_2_. FTE-MYC (FT33-Tag-MYC) cells were a generous gift from Dr. Ronny Drapkin (75). OvTrpMyc-F318LOV cells were a generous gift from Dr. Joe Delaney (76). All reagents were purchased from Fisher Scientific (Hampton, NH) unless otherwise noted. All siRNA was purchased from Bioneer (Oakland, CA) and sequences are listed in Supplemental Table 1. Volasertib was purchased from Cayman Chemical (Ann Arbor, MI).

### Immunoblotting and Immunoprecipitation

Immunoblotting and co-immunoprecipitation was performed as we have previously described (77). Antibodies for immunoblotting and immunoprecipitation included MYC (CST #13987S; RRID: AB_2631168), HSF1 (CST #12972; RRID: AB_2798072), β-actin (CST #3700; RRID: AB_2242334), GAPDH (CST #2118; RRID: AB_561053), and p-HSF1 (S326) (Abcam #76076; RRID: AB_1310328).

### CUT&RUN Sequencing

CUTANA CUT&RUN kit (EpiCypher) was used as previously described (55). Briefly, proliferating OVCAR8 cells were crosslinked with 1% formaldehyde in PBS for 1 minute on culture plates following by glycine quenching. Cells were then scraped and counted with 5e5 cells and then incubated with IgG (CST #3900S; RRID: AB_1550038), MYC (CST #13987S; RRID: AB_2631168), or HSF1 (CST #12972; RRID: AB_2798072) antibodies according to the kit manufacturer’s instructions. Cross-link reversal was performed using 0.8 μL of 10% SDS and 1 μL of 10 μg/μL Proteinase K and overnight incubation at 55°C. DNA was then purified for library generation and next-generation sequencing by the the Center for Genomics and Bioinformatics (CGB) at Indiana University. Libraries were prepared by NEBNext Ultra II DNA Library Preparation Kit protocol (NEB #E7645L) and analyzed by Agilent 4200 TapeStation. The libraries were pooled and loaded on a NextSeq 1000/2000 P2 Reagents (100 cycles) v3 flow cell (#20046811) configured to generated paired end reads. The demultiplexing of the reads was performed using bcl2fastq, version 2.20.0. Raw data was trimmed and aligned to the human (GRCh38/hg38) reference genome using Bowtie2 (78). CUT&RUN peaks were called using MACS2 (79) with peak stringency set to 10^-4^ for HSF1 peaks and 10^-10^ for MYC peaks. Motif enrichment analysis was performed using MEME suite AME (80) and gene binding tracks were visualized using Gviz (81). Gene ontology was performed using Metascape (82).

### RT-qPCR

Total RNA was isolated using the Micro Total RNA isolation kit (Zymo). RNA was reverse transcribed using random primers from a reverse transcription kit (Applied Biosystems). Quantitative PCR was performed using SYBR green master mix (Applied Biosystems) along with gene specific primers using a QuantStudio 3 (Applied Biosystems). Primers used are listed in Supplemental Table 1.

### Luciferase Reporter Assays

The HSF1 activity reporter contains multiple heat shock element (HSE) motifs driving firefly luciferase (Promega #E3751). Experiments were performed with co-transfection of a constitutively active renilla reporter (Addgene #27163; RRID: Addgene_27163). Analysis was completed by dividing firefly activity by renilla activity.

### Immunohistochemistry

Tissues were subjected to immunohistochemistry as we previously described (12). Tissues were purchased commercially from TissueArray.com (#OV8010a). Briefly, slides were deparaffinized and rehydrated prior to antigen retrieval using heat and pressure (2,100 Antigen Retriever; Aptum Biologics). Slides had endogenous peroxidase activity blocked with Bloxall (Vector Laboratories #SP-6000–100) and signal developed with DAB (Vector Laboratories #SK-4105). Antibodies used for IHC included MYC (Abcam #ab32072; RRID: AB_731658), HSF1 (CST #4356; RRID: AB_2120258), p-PLK1 (T210) (Abcam #ab155095). Slides were imaged with Motic Easy Scan and analyzed with QuPath (83).

### Cell Viability and Clonogenic Growth

Cell viability assays were performed with Cell Titer Blue (Promega) according to manufacturer’s instructions. Clonogenic growth assays were performed by seeding <1000 cells into 6-well plates and staining with crystal violet after 5-7 days of growth. Colonies were quantified using FIJI.

### Spheroids

Ovarian cancer cells (2000) were seeded into 24-well ultra-low attachment plates (Corning) and grown in serum-free spheroid media as we previously described (84). Spheroids were grown in the presence or absence of volasertib for 7 days. Spheroids were quantified by manual counting.

### Statistical Analysis

All statistical tests were performed as two-tailed tests. For two-group comparisons, students t-test was used. For multiple group comparisons, ANOVA with Tukey’s post-hoc test was used. All laboratory experiments were completed with a minimum of three biological replicates (e.g. qPCR, luciferase assay, etc.).

### Data Availability Statement

Next generation sequencing data generated in this study was deposited at Gene Expression Omnibus (GEO) at [IN PROCESS]. Publicly available datasets used include The Cancer Genome Atlas (TCGA) cancer datasets that were available via the TCGA Data Portal with analyses performed using cBioportal.org. The Cancer Cell Line Encyclopedia (CCLE) was accessed via DepMap.org. All other raw data generated in this study are available upon request from the corresponding author.

## Results

### MYC and HSF1 are Frequently Co-Amplified in Ovarian Cancer

Copy number changes of the *HSF1* gene have previously been observed across tumor types (85,86). Our analysis of *HSF1* copy number across the TCGA cancer cohorts also observed several cancer types for which *HSF1* has significant copy number gain but ovarian cancer patients had the highest frequency of amplification (≥ 2 copy number gain) (Fig. 1A). The *HSF1* gene is located on chromosome 8q (chr8q) that also contains the *MYC* gene, which also had substantial copy number gains across tumor types with the most frequent gains also in ovarian cancer (Fig. 1B). Analysis of both *MYC* and *HSF1* copy number indicated *MYC* had amplification in 11% of human cancers (1132/9950) while *HSF1* was amplified in 8% of cancers (840/9950) with overlap in a substantial number of patients (Fig. 1C). In fact, greater than 7% of human cancers (730/9950) had co-amplification of both *HSF1* and *MYC* genes (Fig. 1D-E).

**Figure 1:**
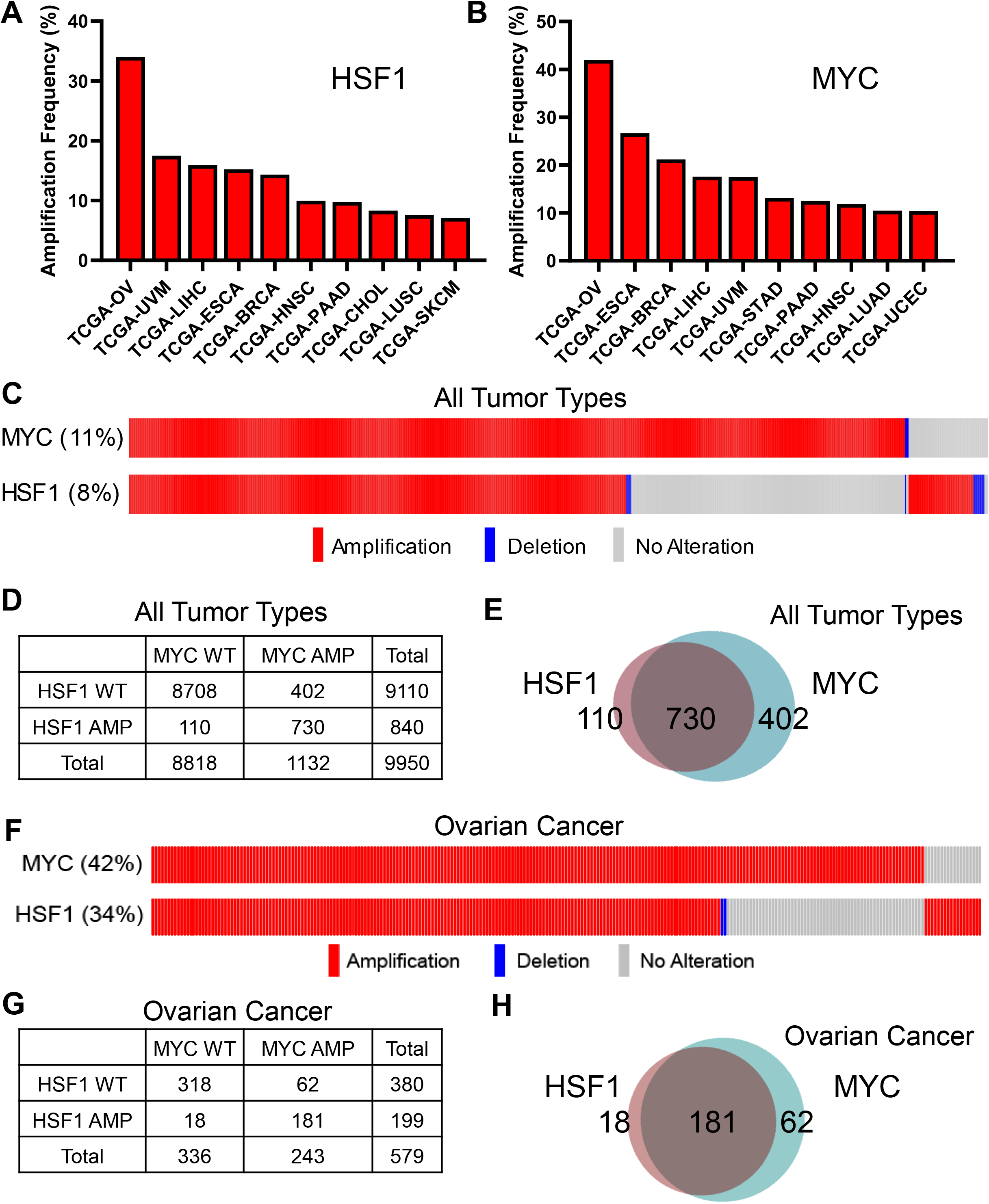
MYC and HSF1 are frequently co-amplified in ovarian cancer. Analysis of copy number variation (CNV) for HSF1 and MYC across cancer types using the TCGA cohorts. Data was analyzed via cBioPortal. A-B) Amplification frequency for HSF1 (A) and MYC (B) across tumor types is presented with amplification defined as ≥2 copy gains. C) Oncoprint for HSF1 and MYC across all TCGA tumor types to indicate overlapping amplification for both genes. D) Chi-square table for HSF1 and MYC amplification across all tumor types. E) Venn diagram of HSF1 and MYC amplification for all tumor types indicating how many amplifications were overlapping. F) Oncoprint for HSF1 and MYC in the TCGA ovarian cancer cohort to indicate overlapping amplifications in ovarian cancer. G) Chi-square table for HSF1 and MYC amplification in ovarian cancer. H) Venn diagram of HSF1 and MYC amplification in ovarian cancer indicating how many amplifications were overlapping. WT=wild type; AMP=amplification.

Considering that both *HSF1* and *MYC* are located on chromosome 8q, their co-amplification could be a random passenger amplification if located on the same amplicon. Our analysis across tumor types showed *MYC* amplification did occur without amplification of *HSF1* in a lower percentage of patients, while *HSF1* was also amplified without *MYC* amplification (Fig. 1D-E). These results suggest that *HSF1* and *MYC* can be separately amplified and not necessarily on the same amplicon. The frequency with which MYC and HSF1 are co-amplified further indicates that co-amplification provides a growth or survival advantage for tumor cells, particularly for ovarian cancer compared to other tumor types, based on the high amplification frequency for HSF1 and MYC in 34% (199/579) and 42% (243/579) of patients, respectively (Fig. 1F-H). There were ovarian cancer patients with single amplification of HSF1 or MYC, but their co-amplification was statistically significant for co-occurrence (Chi-square p<0.0001) (Fig. 1G). Both HSF1 and MYC mRNA expression were strongly associated with gene copy number where mRNA levels for both were highest in tumors with amplifications of each gene (Suppl. Fig. 1A-B), likely indicating a functional role. Interestingly, HSF1 mRNA levels were also elevated with *MYC* amplification but MYC mRNA levels were unchanged with *HSF1* copy number changes (Suppl. Fig. 1C-D). These results suggest that expression of *HSF1* and *MYC* is tied to their own gene copy number but not necessarily tied to copy number changes for other 8q genes, supporting these genes may reside on the same chromosome but on separate amplicons.

### Transcriptional Activity of MYC and HSF1 are Correlated

Because MYC and HSF1 have a high frequency of gene amplification in ovarian cancer, it was of interest to next investigate whether their expression is correlated. By examining the TCGA-OV cohort and ovarian cancer cell lines from the Cancer Cell Line Encyclopedia (CCLE), expression of MYC was weakly associated with expression of HSF1 with correlation coefficients ranging from 0.16-0.20 and not reaching statistical significance in the CCLE (Fig. 2A-B). To next assess their transcriptional activity, we utilized published gene signatures for both MYC (87) and HSF1 (12), which are gene sets downstream of each transcription factor and whose expression is a reflection of MYC and HSF1 transcriptional activity. We found a strong positive correlation between MYC and HSF1 activity in TCGA-OV patients and ovarian cancer cell lines from the CCLE (Fig. 2C-D). We next assessed a panel of ovarian cancer cell lines and found the amplification status of MYC and HSF1 was not perfectly predictive for their protein levels (Fig. 2E). We found no significant differences in MYC or HSF1 transcriptional activities based on copy number for either gene (Suppl. Fig. 2A-D), suggesting that a copy number gain does not necessarily correlate with increased transcriptional activity despite gains in copy number affecting mRNA levels of each gene (Suppl. Fig. 2A-B). For both HSF1 and MYC, expression correlated weakly to HSF1 activity in ovarian cancer (correlation coefficient 0.18-0.27) (Suppl. Fig. 2E-F). The correlation between HSF1 and MYC transcriptional activity is likely to be functional since HSF1 and MYC copy number gains did not correspond with transcriptional activity.

**Figure 2:**
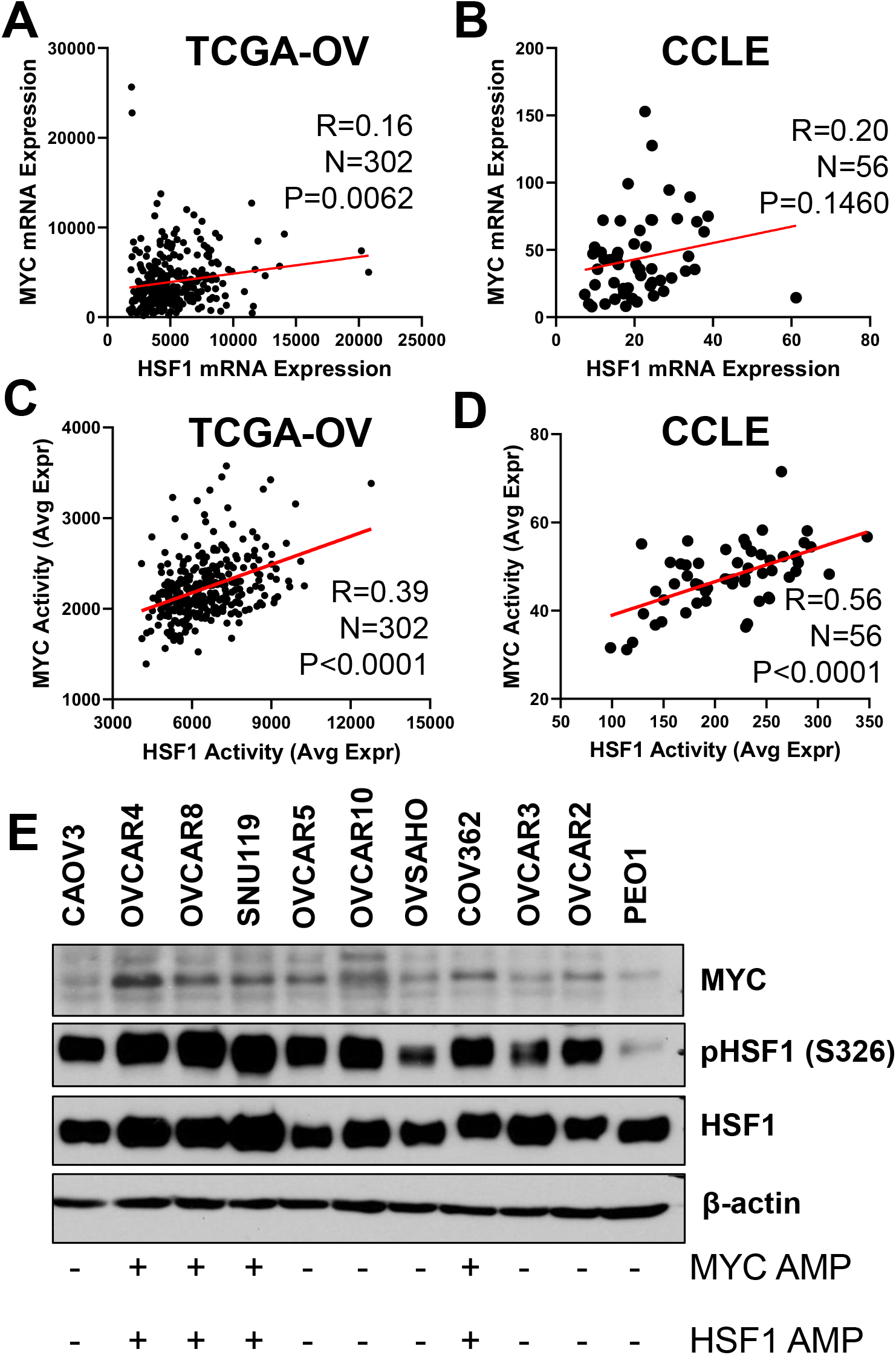
HSF1 activity is associated with MYC activity. A-B) HSF1 mRNA levels are plotted with MYC mRNA levels and analyzed with Pearson correlation in the TCGA-OV cohort (A) and ovarian cancer cell lines from the cancer cell line encyclopedia (CCLE) (B). C-D) HSF1 and MYC transcriptional activities were calculated using published gene signatures and subjected to Pearson correlation using the TCGA-OV cohort (C) and ovarian cancer cell lines from the CCLE (D). E) Total protein from indicated cell lines were subjected to immunoblotting with the indicated antibodies. Immunoblotting was performed with three independent replicates.

### Overlapping MYC and HSF1 Gene Targets in Ovarian Cancer Cells

Considering the strong correlation between MYC and HSF1 transcriptional activity, we investigated whether they share similar gene targets, as suggested by a recent report on reduced global MYC DNA binding in the absence of HSF1 (55). We assessed genome binding patterns for both MYC and HSF1 by CUT&RUN sequencing. We selected the OVCAR8 cell line, in which both genes are amplified. For quality control purposes, we compared the CUT&RUN to previously published ChIP-seq data for both MYC (88) and HSF1 (29). There was substantial overlap in the gene targets identified from the current CUT&RUN and these previous ChIP-Seq samples (Suppl. Fig. 3A-D). Furthermore, the binding motifs for MYC and HSF1 were highly enriched in their respective CUT&RUN samples (Suppl. Fig. 3E-F). Lastly, gene ontology for MYC and HSF1 bound genes reported common ontologies associated with these transcription factors (Suppl. Fig. 3G-H). These results demonstrate the binding profiles detected through this CUT&RUN are reflective of true binding patterns for MYC and HSF1.

As expected, MYC binding across the genome was considerable (>12,000 peaks called) while HSF1 had over 2,100 peaks across the genome. These peaks translated to >8,000 unique genes bound by MYC and >1,500 unique genes bound by HSF1. There were 877 peaks that displayed overlap in MYC and HSF1 binding, which represented 39% of all HSF1 binding peaks but less than 7% of all MYC peaks due to the large number of MYC peaks detected (Fig. 3A). There were 760 unique genes that had overlapping peaks for both MYC and HSF1 representing ∼9% of MYC bound genes but 44% of genes bound by HSF1 (Fig. 3B). There were another 392 genes wherein both MYC and HSF1 were bound but did not have overlapping peaks. Consistent with these results, the MYC binding motif was significantly enriched in the HSF1 CUT&RUN and the HSF1 binding motif was significantly enriched in the MYC CUT&RUN (Fig. 3C-D). MYC bound peaks were largely at intergenic regions, introns, and promoters with a lower percentage of bound sites located on exons (Fig. 3E). HSF1 bound peaks showed more binding to introns and intergenic regions with a lower number of bound sites at promoters (Fig. 3F). Analyzing only the MYC and HSF1 overlapping peaks resulted a more equal distribution of MYC and HSF1 binding across promoters, introns, and intergenic regions (Fig. 3G). We observed variations in binding between MYC and HSF1 across the genome and found genes that are bound only by MYC (CDK4, ODC1), genes that are bound only by HSF1 (PROM2, RBM23), and genes that are bound by both MYC and HSF1 (HSPA1B, JMJD6) (Fig. 3H).

**Figure 3:**
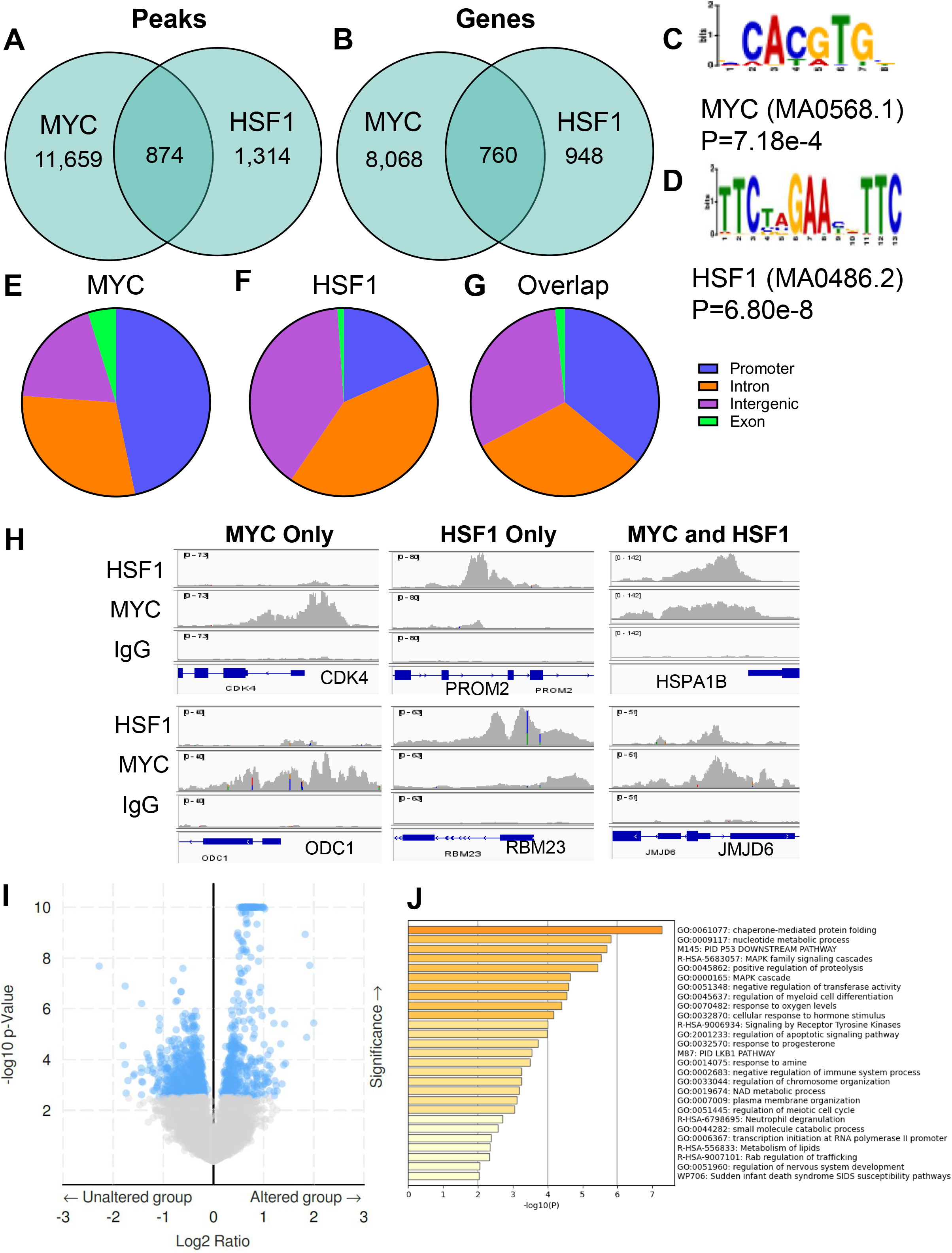
HSF1 and MYC share binding locations in the genome of ovarian cancer cells. OVCAR8 cells were subjected to CUT&RUN for both HSF1 and MYC. A-B) HSF1 and MYC called peaks (A) and annotated genes (B) are presented with the overlapping number between HSF1 and MYC in the Venn diagram overlap. C-D) Motif analysis showing the MYC binding motif that presented in the HSF1 CUT&RUN (C) and the HSF1 binding motif that presented in the MYC CUT&RUN (D). E-G) Binding peaks for MYC (E), HSF1 (F), and the genes comprising the overlapping peaks (G) were classified into their binding location. H) Gene tracks from HSF1 and MYC CUT&RUN showing examples of genes with MYC only binding (left), HSF1 only binding (middle), or genes with binding of both MYC and HSF1 (right). I) Volcano plot showing differentially expressed genes from the TCGA-OV cohort comparing tumors with MYC-HSF1 co-amplification (altered group) vs. tumors with MYC-HSF1 WT copy number (unaltered group). J) Genes that were significantly higher expressed in MYC-HSF1 co-amplified tumors from (I) were overlapped with genes were bound by both MYC and HSF1 (n=83). These upregulated and MYC-HSF1 bound genes were subjected to gene ontology and enriched categories are presented.

To ascertain which of these MYC-HSF1 bound genes may be functional in ovarian cancer, differential gene expression analysis was performed on the TCGA-OV cohort between samples with or without MYC-HSF1 co-amplification. We identified 4,155 differentially expressed genes between these two groups with 1,480 genes higher expressed in MYC-HSF1 co-amplified tumors and 2,675 genes higher expressed in tumors that have WT copy number of MYC and HSF1 (Fig. 3I). Of the 1,480 genes that were upregulated in MYC-HSF1 co-amplified tumors, 83 of those genes had overlapping MYC and HSF1 peaks. Gene ontology was performed using these 83 genes that resulted in some expected enriched categories with clear association to MYC and/or HSF1 functions such as chaperone-mediated folding, metabolism, and transcription (Fig. 3J). In addition, there were several other enriched categories that support tumor development and progression such as MAPK signaling, response to oxygen levels, and receptor tyrosine kinase signaling among others (Fig. 3J). These data suggest that amplified MYC and HSF1 are activating a program to support the initiation and growth of tumor cells by upregulating growth-promoting pathways, metabolism and oxygen-sensing pathways to meet these increased energy needs, and protein-folding machinery to protect against imbalances in proteostasis.

### HSF1 and MYC Functionally Interact in Ovarian Cancer Cells

Analysis of the HSF1 and MYC CUT&RUN identified MYC binding to the *HSF1* promoter as well as both HSF1 and MYC bound to the *MYC* promoter in OVCAR8 cells (Fig. 4A). Knockdown of MYC lowered levels of HSF1 protein but knockdown of HSF1 had little effect on MYC protein levels in OVCAR8 cells (Fig. 4B-C). Loss of HSF1 had no effect on MYC in human fallopian tube epithelial cells (FTEs), the cell of origin of HGSOC, that had been engineered to overexpress MYC (75) (Fig. 4D). These data suggest MYC has a larger effect on HSF1 expression than the reverse. This was confirmed at the RNA level where knockdown of HSF1 had minimal effect on MYC RNA levels while knockdown of MYC resulted in a modest, but significant, decrease in HSF1 expression (Fig. 4E-F). Conversely, overexpression of MYC did not affect HSF1 expression or vice versa (Fig. 4G-H), indicating HSF1 expression may be partially dependent on MYC but overexpression of MYC is not sufficient to drive further expression of HSF1. Consistent with these results, MYC activity in the TCGA-OV cohort was significantly associated with HSF1 expression (Suppl. Fig. 4A) but HSF1 activity was not associated with MYC expression (Suppl. Fig. 4B).

**Figure 4:**
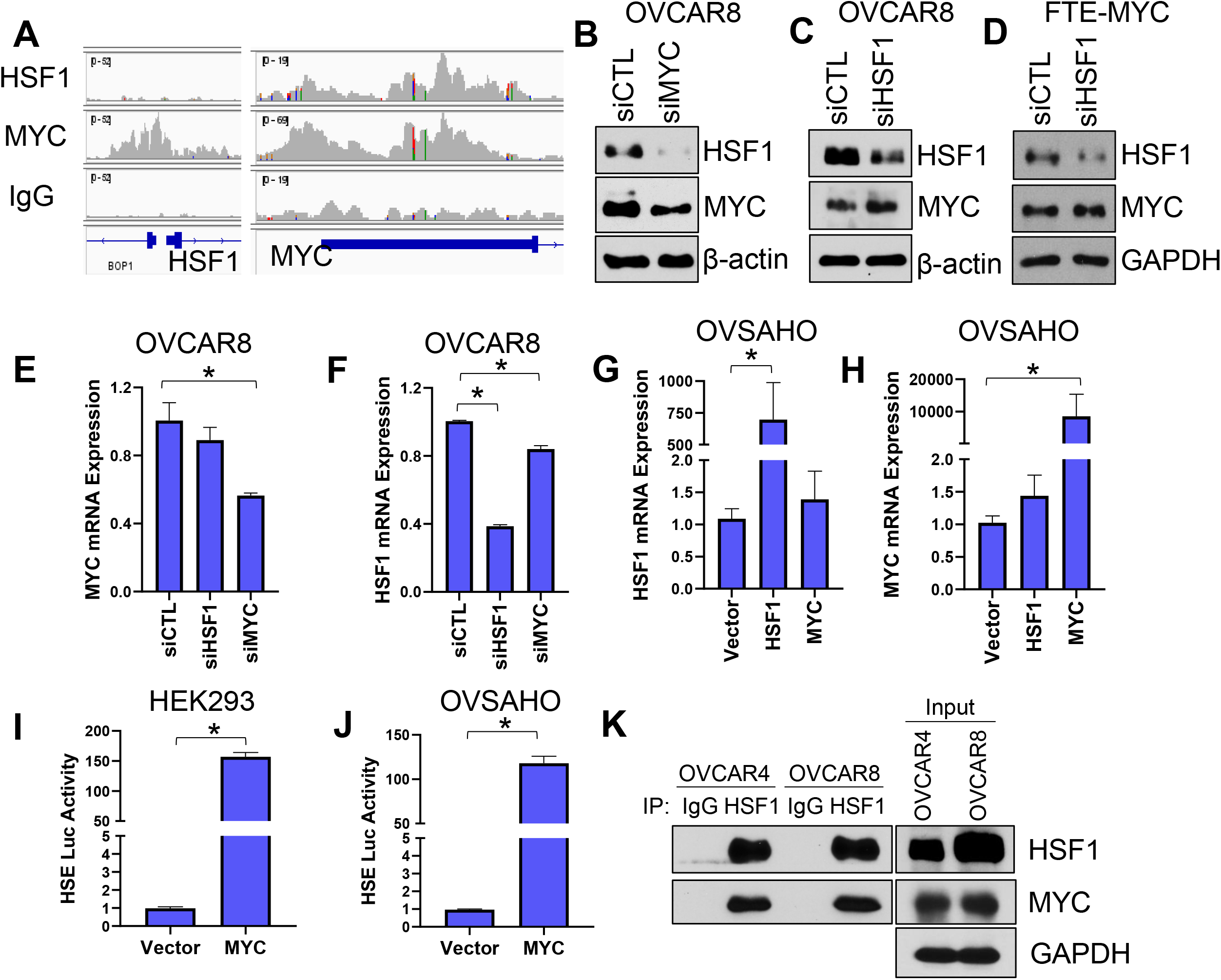
HSF1 and MYC cooperate in ovarian cancer cells. A) Analysis of HSF1 and MYC CUT&RUN showing gene tracks and binding at the MYC and HSF1 genes. B-C) OVCAR8 cells were transfected with MYC (B) or HSF1 (C) siRNA for 48 hrs. Total protein was subjected to immunoblotting. D) FTE-MYC were transfected with HSF1 siRNA for 48 hrs. Total protein was subjected to immunoblotting. E-F) OVCAR8 cells were transfected with control (CTL), HSF1, or MYC siRNA for 48 hrs. Total RNA was subjected to RT-qPCR for MYC (E) and HSF1 (F). G-H) OVSAHO cells were transfected with vector, HSF1 or MYC for 48 hrs. Total RNA was subjected to RT-qPCR for HSF1 (G) and MYC (H). I-J) HEK293 (I) and OVSAHO (J) cells were transfected with pGL4-HSE HSF1 luciferase reporter along with vector or MYC. Luciferase activity was measured by luminescence in the presence of d-luciferin and normalized to a control renilla activity. K) Total protein from OVCAR4 and OVCAR8 cells were subjected to immunoprecipitation with HSF1 antibodies and blotted with indicated antibodies. *Indicates p<0.05.

It has been previously shown that HSF1 can potentiate MYC transcriptional activity (55) but MYC can affect HSF1 transcriptional activity has not been tested. Overexpression of MYC in non-cancer HEK293 cells (Fig. 4I) or HGSOC OVSAHO cells (Fig. 4J) resulted in a significantly increased activation of a heat shock element (HSE) luciferase reporter, likely due to an effect of MYC on HSF1 expression but could possibly be due to a cooperation between the HSF1 and MYC proteins (55). Protein level cooperation between HSF1 and MYC would also be consistent with overlapping bound regions by HSF1 and MYC (Fig. 3). In HGSOC cell lines with both genes amplified, we detected a physical interaction between HSF1 and MYC in ovarian cancer cell lines that have these two genes amplified (Fig. 4J). These data suggest a physical interaction and cooperation between MYC and HSF1 that ultimately supports their function to promote ovarian cancer cell growth.

### HSF1-MYC Co-Amplification drive HGSOC Cell Growth

It was next of interest to investigate the effect of HSF1 and/or MYC amplification on HGSOC cell growth. Loss of either HSF1 or MYC in OVCAR8 cells significantly reduced clonogenic growth (Fig. 5A) and cell proliferation (Fig. 5B). Similarly, loss of HSF1 in FTE-MYC cells (75) significantly reduced clonogenic growth (Fig. 5C) and proliferation (Fig. 5D). We further tested the dependency of FTE-MYC cells and OvTrpMyc-FOV318, which are MYC and mutant p53-driven mouse HGSOC cells on HSF1 using the HSF1-specific inhibitor SISU-102 (DTHIB) (89). Treatment with SISU-102 inhibited clonogenic growth of both FTE-MYC and OVTrpMyc-FOV318 cells (Fig. 5E-F). These data support a model whereby HSF1 and MYC can cooperate to drive cell growth and dependency on both transcription factors.

**Figure 5:**
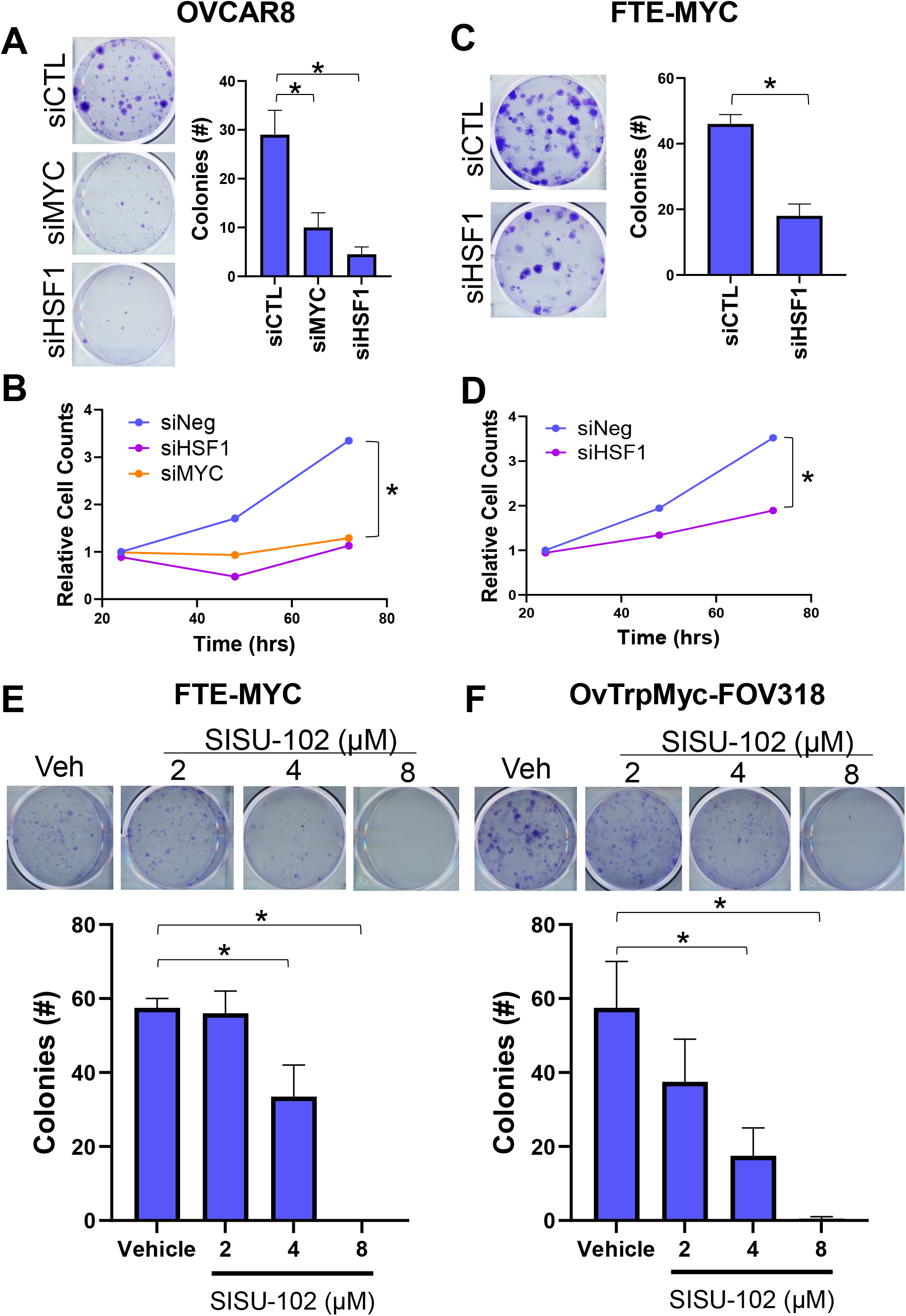
HSF1-MYC Co-amplified cells require both HSF1 and MYC for growth. A-B) OVCAR8 cells were transfected with control (CTL), HSF1, or MYC siRNA for 48 hrs followed by a clonogenic growth assay for 7 days (A) or cell proliferation (B). C-D) FTE-MYC cells were transfected with control (CTL) or HSF1 siRNA for 48 hrs followed by a clonogenic growth assay for 7 days (C) or cell proliferation (D). E-F) Either FTE-MYC (E) or OvTrpMyc-FOV318 cells (F) were subjected to clonogenic growth assay for 7 days in the presence of vehicle or the HSF1 inhibitor SISU-102 at the indicated dosages.

### MYC and HSF1 are Associated with PLK1 in Human High Grade Serous Ovarian Cancer

To identify possible therapeutic approaches that would benefit HGSOC tumors with MYC and HSF1 co-amplification, polo-like kinase 1 (PLK1) was identified as a kinase that can directly and indirectly regulate both MYC and HSF1 (69–73). MYC has also previously been shown to bind and upregulate the PLK1 gene (90), which we also observed in OVCAR8 cells (Suppl. Fig. 5A). Consequently, PLK1 could be an attractive therapeutic target to abolish this cancer-promoting signaling node (Fig. 6A). We first assessed the correlation of MYC and HSF1 activity with PLK1 activity in the TCGA-OV cohort using a published gene signature for PLK1 (91). PLK1 activity was significantly associated with both MYC and HSF1 activity (Fig. 6B-C). To further assess whether active PLK1 was associated with MYC or HSF1 in primary ovarian cancer patient samples, we performed IHC for active PLK1 (pT210), HSF1, and MYC using 58 high grade serous ovarian cancer patient tumors (Fig. 6D). Nuclear levels of MYC and HSF1 were correlated (Fig. 6E), confirming activity of the two transcription factors in the same tumors. Active PLK1 correlated with nuclear levels of both HSF1 and MYC (Fig. 6F-G), indicating a positive association between active PLK1, MYC and HSF1. These observations support that PLK1 could be an attractive therapeutic target to abolish a cancer-promoting signaling node.

**Figure 6:**
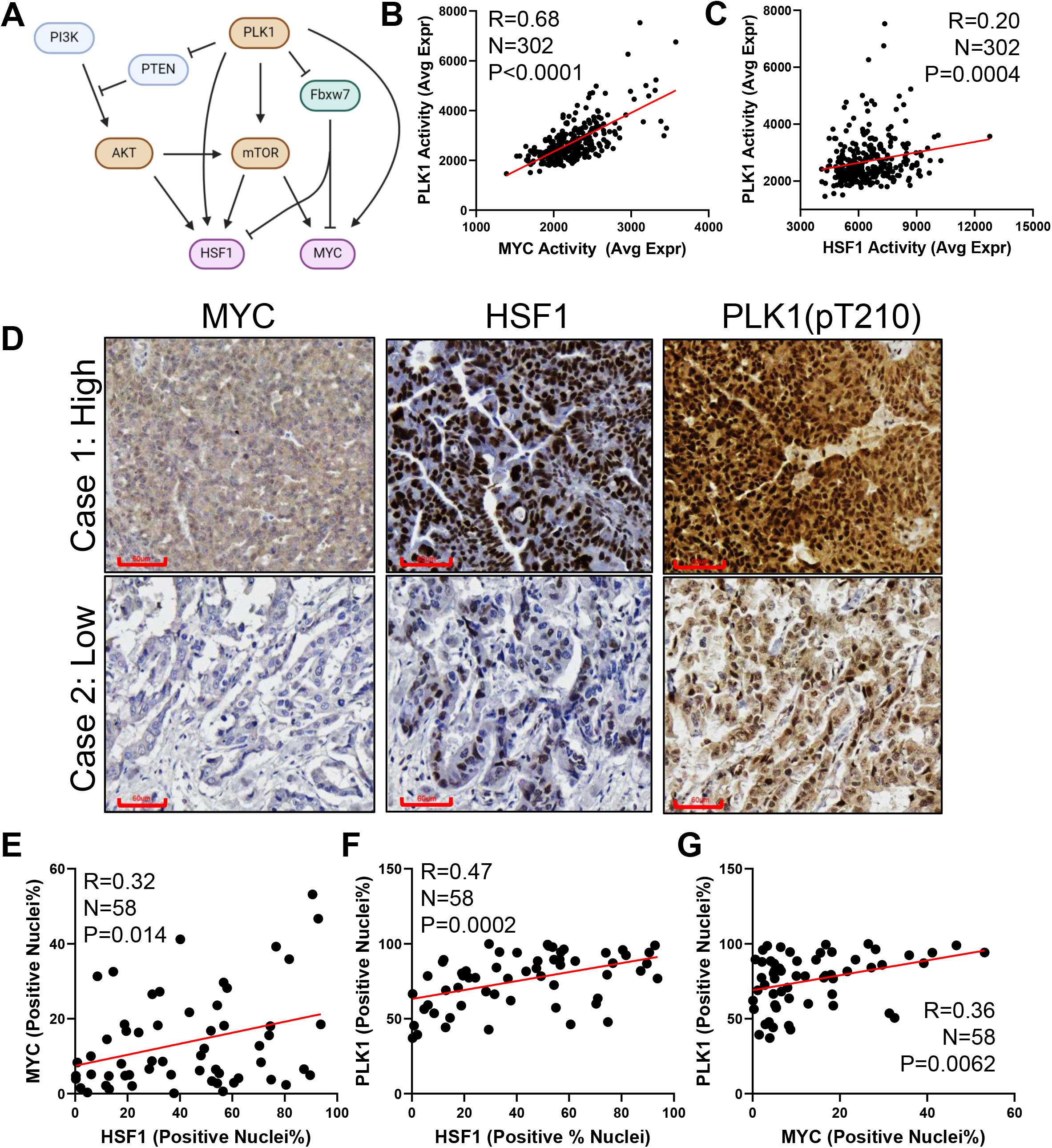
HSF1 and MYC are correlated with active PLK1 in ovarian tumors. A) Diagram of the relationship between PLK1 and MYC/HSF1 indicating PLK1 can directly regulate MYC and HSF1 through phosphorylation but also indirectly by regulating the PI3K-AKT-mTOR and FBXW7 pathways. B-C) PLK1 activity was assessed using a published gene signature in the TCGA-OV cohort and correlated with activity signatures for MYC (B) or HSF1 (C). D) A cohort of 58 high grade serous ovarian tumors were subjected to immunohistochemistry with antibodies for MYC, HSF1, and active PLK1 (pT210). E-F) Immunohistochemistry in D was analyzed with quPath to identify percent of cells that have positive nuclei for these markers to indicate active levels. Pearson correlation was used to correlate active HSF1 and MYC (E), HSF1 and PLK1 (F), and MYC and PLK1 (G).

### PLK1 Inhibition is More Effective with MYC and HSF1 Dual Amplification

PLK1 is an active therapeutic target with several compounds targeting this kinase in clinical trials. In particular, volasertib (BI-6727) is a selective PLK1 inhibitor that has shown promise as an effective cancer therapy in early phase clinical trials (92–94). To assess whether volasertib has specificity for MYC-HSF1 dual-amplified ovarian cancer cells, we first determined the IC_50_ for volasertib in cell lines with or without HSF1-MYC co-amplification. The average IC_50_ for volasertib in HSF1-MYC co-amplified cell lines was 33 nM whereas cells that were wild-type (WT) for HSF1 and MYC had an average IC_50_ of 7.8 μM, for a >200-fold difference (Fig. 7A). Volasertib also showed a greater reduction in clonogenic growth for MYC-HSF1 co-amplified cells compared to MYC-HSF1 WT cells (Fig. 7B-C). Consistent with genetic depletion or inhibition of HSF1, volasertib also significantly reduced clonogenic growth of FTE-MYC cells (Fig. 7D). Furthermore, volasertib significantly reduced the ability of HSF1-MYC co-amplified OVCAR8 cells to form spheroids under low attachment conditions (Fig. 7E), while spheroid growth of MYC-HSF1 WT CAOV3 cells was not significantly altered (Fig. 7E).

**Figure 7:**
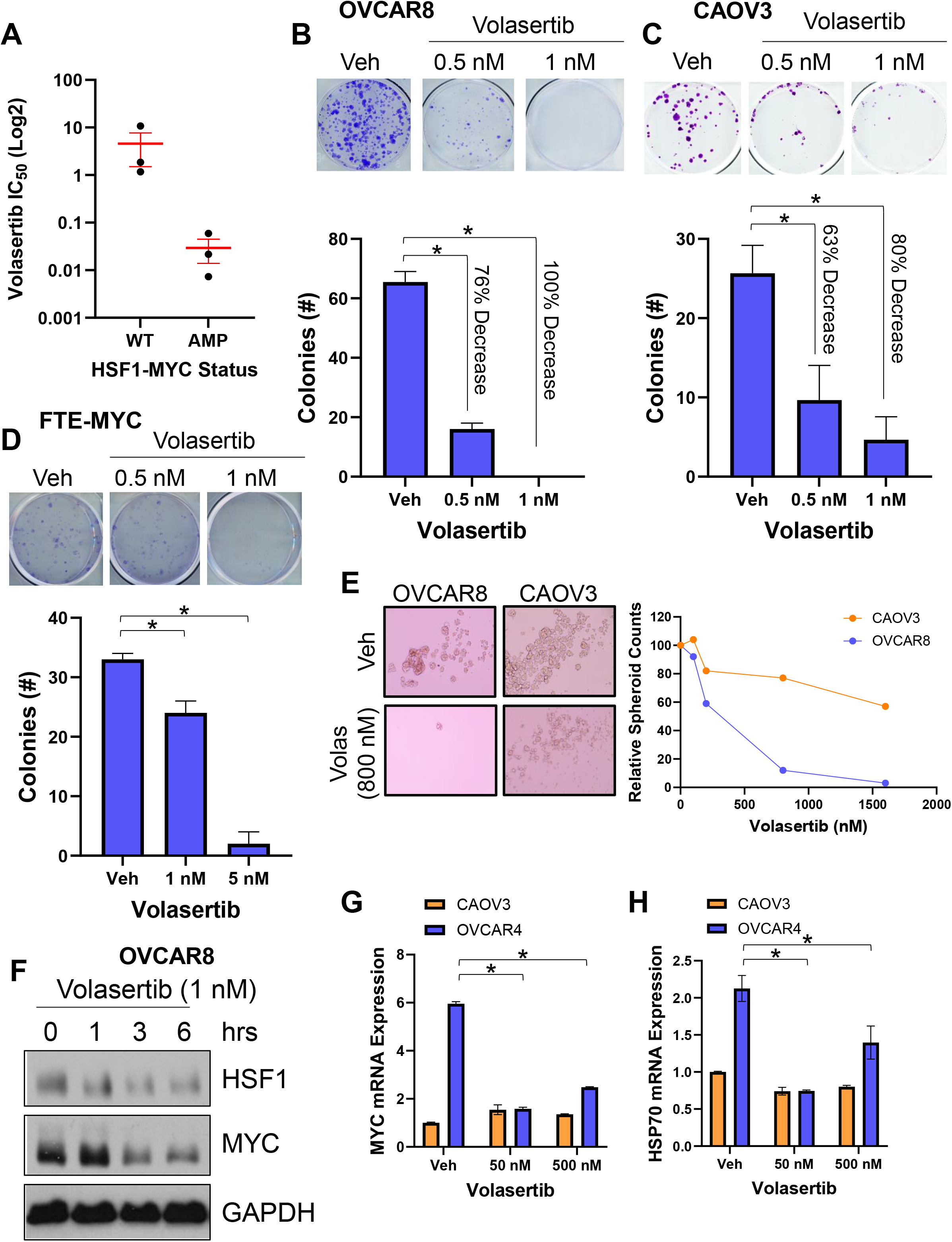
HSF1-MYC co-amplified cells are highly sensitive to PLK1 inhibition with volasertib. A) IC_50_ for volasertib in human ovarian cancer cells with HSF1 and MYC amplified or WT. B-D) OVCAR8 (B, AMP), CAOV3 (C, WT), or FTE-MYC (D) cells were subjected to a clonogenic growth assay for 7 days in the presence of vehicle or volasertib at indicated doses. E) OVCAR8 and CAOV3 cells were subjected to tumor spheroid growth for 12 days in the presence of vehicle or volasertib at the indicated doses. F) OVCAR8 cells were grown in the presence of vehicle or volasertib (1 nM) for the indicated time periods. Total protein was subjected to immunoblotting for the indicated antibodies. G-H) OVCAR4 (AMP) or CAOV3 (WT) cells were treated with vehicle or volasertib at the indicated doses for 24 hrs. Total RNA was subjected to RT-qPCR for MYC (G) or HSP70 (H).

As PLK1 phosphorylation has been reported to increase the protein stability of MYC and HSF1 (69–73), we tested whether volasertib affects MYC or HSF1 protein levels in co-amplified OVCAR8 cells. A time course of volasertib exposure indicated a loss of MYC and HSF1 protein within 1-3 hours of 1 nM volasertib exposure (Fig 7F), suggesting volasertib is suppressing PLK1-mediated protein stabilization of HSF1 and MYC. Similarly, volasertib was also found to suppress expression of MYC and HSP70, a direct target of HSF1, in MYC-HSF1 co-amplified OVCAR4 cells but not in CAOV3 cells that have WT MYC and HSF1 copy number. Taken together, inhibition of PLK1 can destabilize the HSF1 and MYC signaling node in ovarian cancer cells, indicating HSF1-MYC co-amplification could serve as a potential biomarker for therapeutic response to PLK1 inhibition.

## Discussion

Ovarian cancer is a deadly cancer in women that initially responds to platinum-based therapies but is almost universally recurrent with the development of platinum resistance. Platinum-based therapy regimens are ubiquitously used in ovarian cancer patients regardless of the underlying molecular nature of the tumor. Here we describe a biomarker related to increased copy number on chromosome 8 leading to gene amplification of both HSF1 and MYC, which we demonstrate can be a marker for sensitivity to PLK1 inhibition. While PLK1 inhibitors have not met expectations in clinical trials, these results would suggest targeting PLK1 using a precision medicine approach would have a higher likelihood of success with HSF1-MYC co-amplification as one potential biomarker. These data demonstrate that HSF1-MYC co-amplification is present in a number of different cancer types but with specific enrichment in ovarian cancer, for reasons unknown at this point. These data imply HSF1-MYC co-amplification could be a biomarker applicable to a large number of cancer patients and approximately one-third of ovarian cancer patients. Therefore, PLK1 inhibitors may be able to find a place in treating patients with this co-amplification, specifically in this large percentage of ovarian cancer patients, perhaps warranting further clinical testing using this biomarker.

PLK1 has been considered a viable therapeutic target in cancer for many years (95–97). Volasertib (BI-6727) is a selective PLK1 inhibitor that has shown promise as an effective cancer therapy in early phase clinical trials with side effects being reversible and manageable (92–94,98,99). However, Phase II trials were disappointing with only modest antitumor activity as a monotherapy (93,100,101). One of these Phase II trials occurred with advanced ovarian cancer patients and showed the median progression-free survival (PFS) for volasertib (13.1 weeks) was worse than chemotherapy (20.6 weeks). However, there were six patients in the study receiving volasertib that achieved a PFS of more than one year while no patients receiving chemotherapy achieved a PFS greater than one year (93). Additionally, patients receiving chemotherapy discontinued treatment to adverse events more frequently than volasertib (93). The study did not perform any evaluation or analyses related to potential biomarkers that could delineate these patients who responded well to volasertib. The current study would suggest that HSF1-MYC co-amplification could potentially serve as a biomarker for patients that would response more favorably to volasertib or other PLK1 inhibitors. While it seems PLK1 inhibition in cancer has lost some favor due to these trials not meeting adequate endpoints, the current study suggests that a precision medicine approach to find patients that would be more likely to respond favorably may be the path forward for PLK1 inhibition in patients.

The current study also adds the ongoing discoveries indicating a biological, and physical, interaction between HSF1 and MYC. MYC is a frequent oncogenic driver across many tumor types. It has previously been reported that MYC-driven hepatocellular carcinoma requires HSF1 for tumor formation (56) and results from the current study support this dependency also occurs in ovarian cancer. Not surprisingly, HSF1 and MYC were also co-amplified in hepatocellular carcinoma as well but in a lower percentage of patients than ovarian cancer. A recent elegant study demonstrated that HSF1 potentiates MYC transcriptional activity driven by a physical interaction between the transcription factors and that HSF1 was essential to recruitment of epigenetic machinery required for gene regulation (55). While these studies were not in cancer cells, the current results support that MYC and HSF1 also form a protein complex in ovarian cancer cells and MYC-driven ovarian cancer cells are dependent on HSF1. The current study also found a significant number of binding sites for HSF1 and MYC in ovarian cancer cells wherein the binding peaks are overlapping, which would be consistent with previous reports suggesting HSF1 and MYC protein complexes form with DNA (55). These and previous results do suggest the MYC and HSF1 interaction will be the subject of future investigation as this interaction is likely to be critical to tumorigenesis and perhaps several other functions known to be driven by MYC and HSF1 such as the cancer stem-like population among many others (36,102).

The current study is the first to show genome binding patterns for both HSF1 and MYC in a co-amplified ovarian cancer cell. These data further support a gene regulating protein complex involving both HSF1 and MYC evidenced by highly similar binding peaks at a large number of gene targets. These data also indicate a MYC-HSF1 complex that can bind both proximal promoters and distal enhancers, consistent with previous observations that these transcription factors can function at both promoters and enhancer locations in cancer cells (103,104). Interestingly, we also found that both the MYC and HSF1 genes are also targets with MYC binding the HSF1 promoter and both MYC and HSF1 binding the MYC promoter. This translated to HSF1 expression having partial dependency on MYC, which may be partial due to the high HSF1 copy number since HSF1 expression was shown to have greater dependency on MYC in non-cancer cells from a previous study (55). Future studies will ascertain the importance of the physical interaction between HSF1 and MYC by map the necessary regions for the interaction in the hopes of blocking their physical interaction without compromising protein levels.

The specific vulnerability of HSF1-MYC co-amplified cells to PLK1 inhibition seems to originate with the signaling node relating HSF1 and MYC to other growth promoting pathways. Aside from directly phosphorylating both MYC and HSF1, PLK1 can also indirectly affect MYC and HSF1 function through phosphorylation and deactivation of PTEN thereby enhancing activity of PI3K-mediated signaling leading to AKT and mTOR activation (69,71,72,105,106), both of which have been shown to activate HSF1 while mTOR can also phosphorylate MYC (39,107,108). PLK1 can also inactivate the E3 ligase Fbxw7 (90), which has been shown to downregulate both MYC and HSF1 (27). FBXW7 additionally suppresses several other growth promoting pathways including Cyclin E and NOTCH1 (109). The current data indicate MYC can also directly bind the PLK1 promoter, possibly indicating this signaling network can reinforce its own activity through a positive feedback cycle that would lead to consistent proliferation, implying it could be critical for tumor initiation. Therefore, these cancer cells are provided a clear growth advantage by activation of this signaling network that enables efficient proliferation and suppression of growth arresting pathways.

The current study supports ongoing publications linking HSF1 and MYC transcription factors (55,56). Along with the current study, the link between HSF1 and MYC has been seen in cancer cells (56) and non-cancer cells (55), suggesting their interaction is physiological but can be exacerbated in cancer cells. MYC has previously been shown to cooperate with the unfolded protein response through ATF4 (110). MYC cooperation with HSF1 may indicate that MYC cooperates with several stress response and proteostasis pathways possibly due to the positive effects of MYC on protein translation. Several attempts to therapeutically target MYC or HSF1 have all largely failed to reach clinical trials. There is currently an ongoing Phase Ia trial for an HSF1 Pathway inhibitor (NXP800) (111) (NCT05226507) that was promoted to a Phase Ib trial that is the first-in-human compound to inhibit HSF1. Future studies to further clarify the interaction of these two important transcriptional pathways will be exciting if they will reveal viable therapeutic targets to disrupt these cancer promoting pathways that could lead to clinical advances in a precision medicine therapeutic approach.

## Acknowledgements

This publication was made possible, in part, with support from the National Cancer Institute (K22CA207575; RLC). Additional funding support was provided by the Ovarian Cancer Research Alliance Mentored Investigator Grant (IW). Further support was provided by the Van Andel Institute through the Van Andel Institute – Stand Up To Cancer Epigenetics Dream Team (RLC, KPN). The indicated Stand Up To Cancer Grant is administered by the American Association for Cancer Research, the Scientific Partner of SU2C. The content is solely the responsibility of the authors and does not necessarily represent the official views of the National Institutes of Health. We would also like to acknowledge the Light Microscopy Imaging Center, the Center for Medical Genomics, and the Center for Genomics and Bioinformatics at Indiana University for use of their core facilities.

---

**Supplemental Figure 1:**
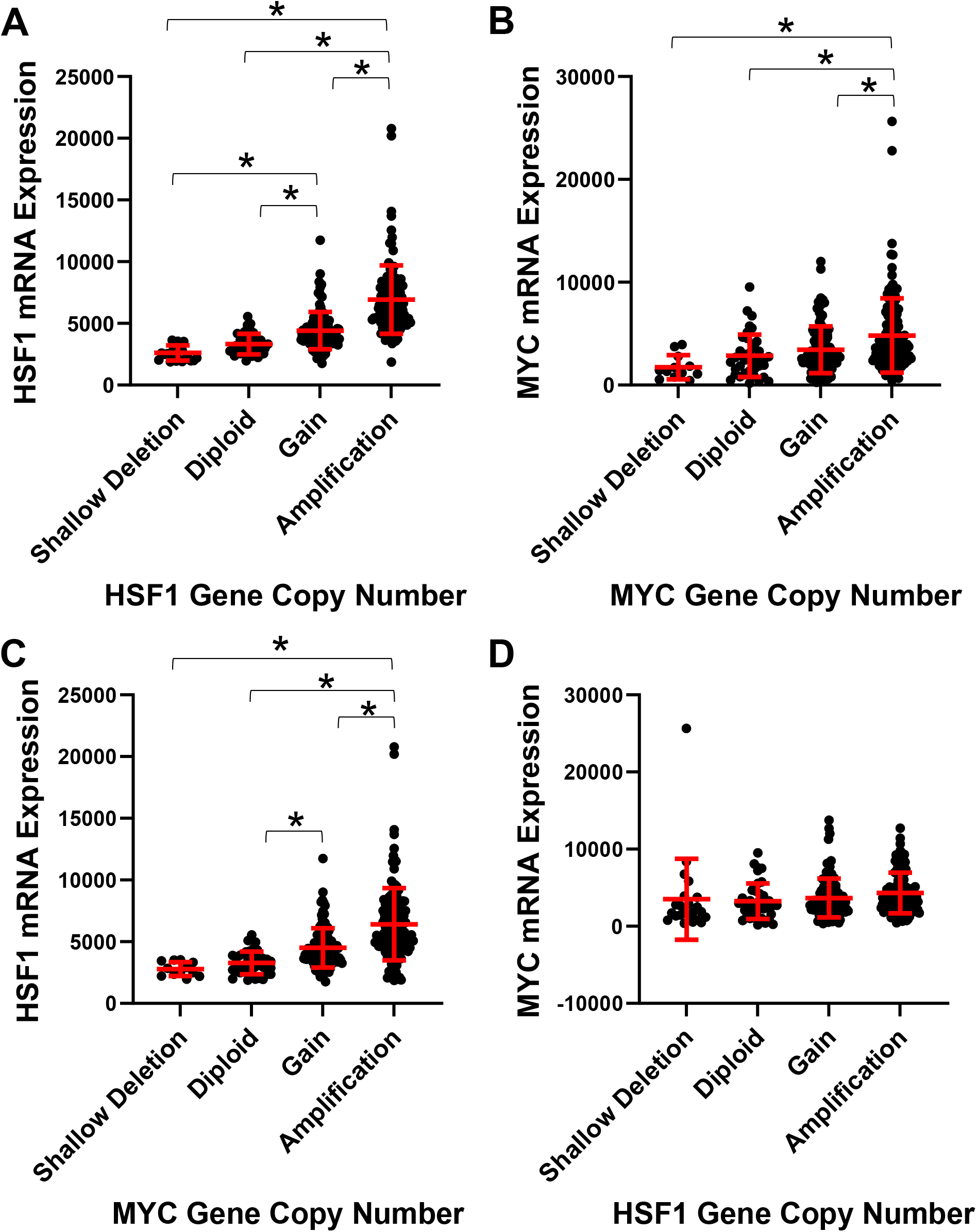
Effect of HSF1 or MYC copy number on mRNA expression. The TCGA-OV cohort was used for this analysis and performed using cBioportal.org. A-B) Expression of HSF1 (A) and MYC (B) are presented in relation to their respective copy number changes. C-D) Expression of HSF1 as it relates to MYC copy number (C) and expression of MYC as it relates to HSF1 copy number. *Indicates p<0.01.

**Supplemental Figure 2:**
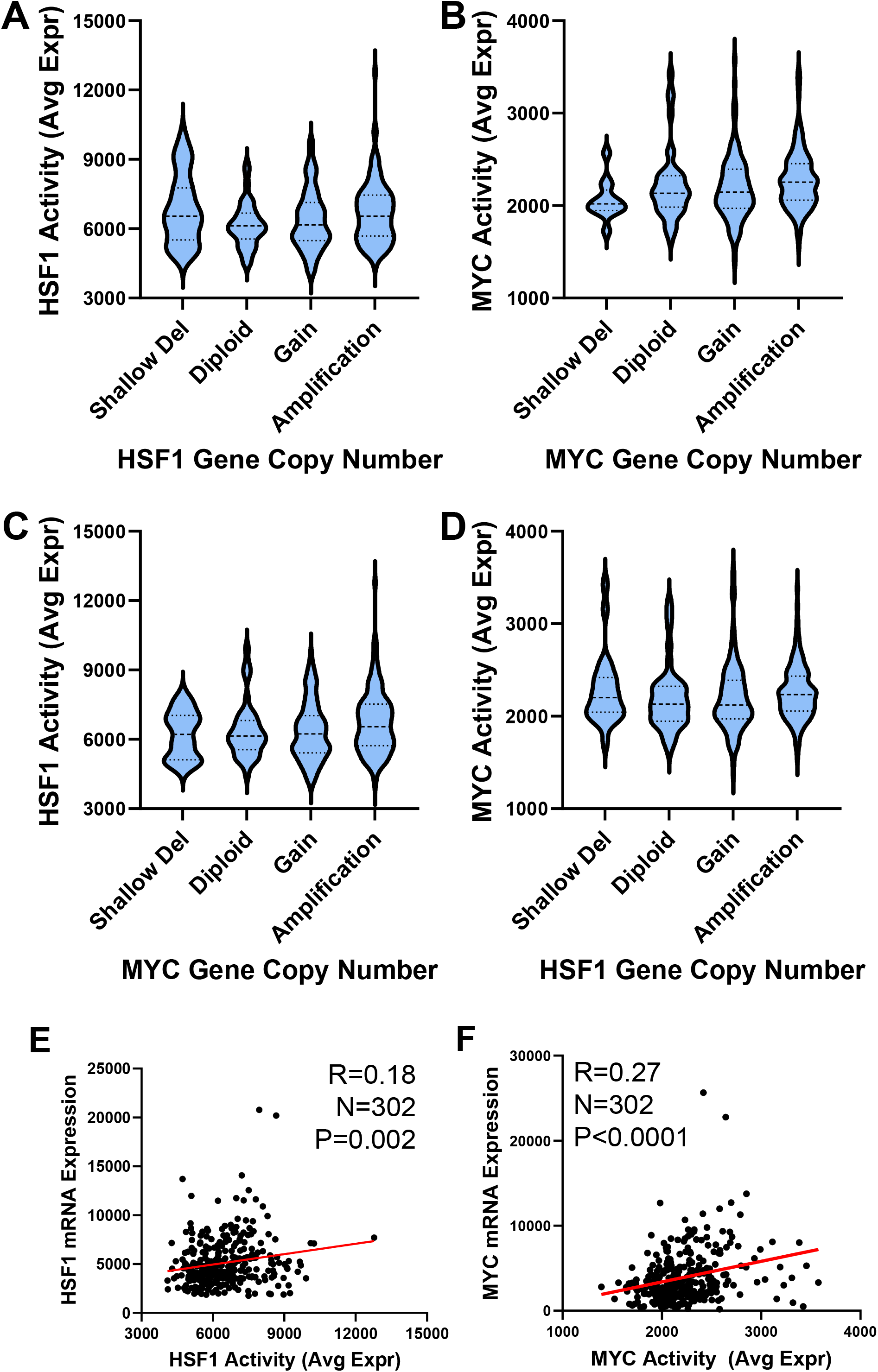
Effect of HSF1 or MYC copy number on their transcriptional activity. MYC and HSF1 activity was calculated using published gene signatures in the TCGA-OV cohort. A-B) HSF1 activity (A) and MYC activity (B) are plotted by gene copy number of each respective gene. C-D) HSF1 activity (C) and MYC activity (D) are plotted by gene copy number of the opposite gene.

**Supplemental Figure 3:**
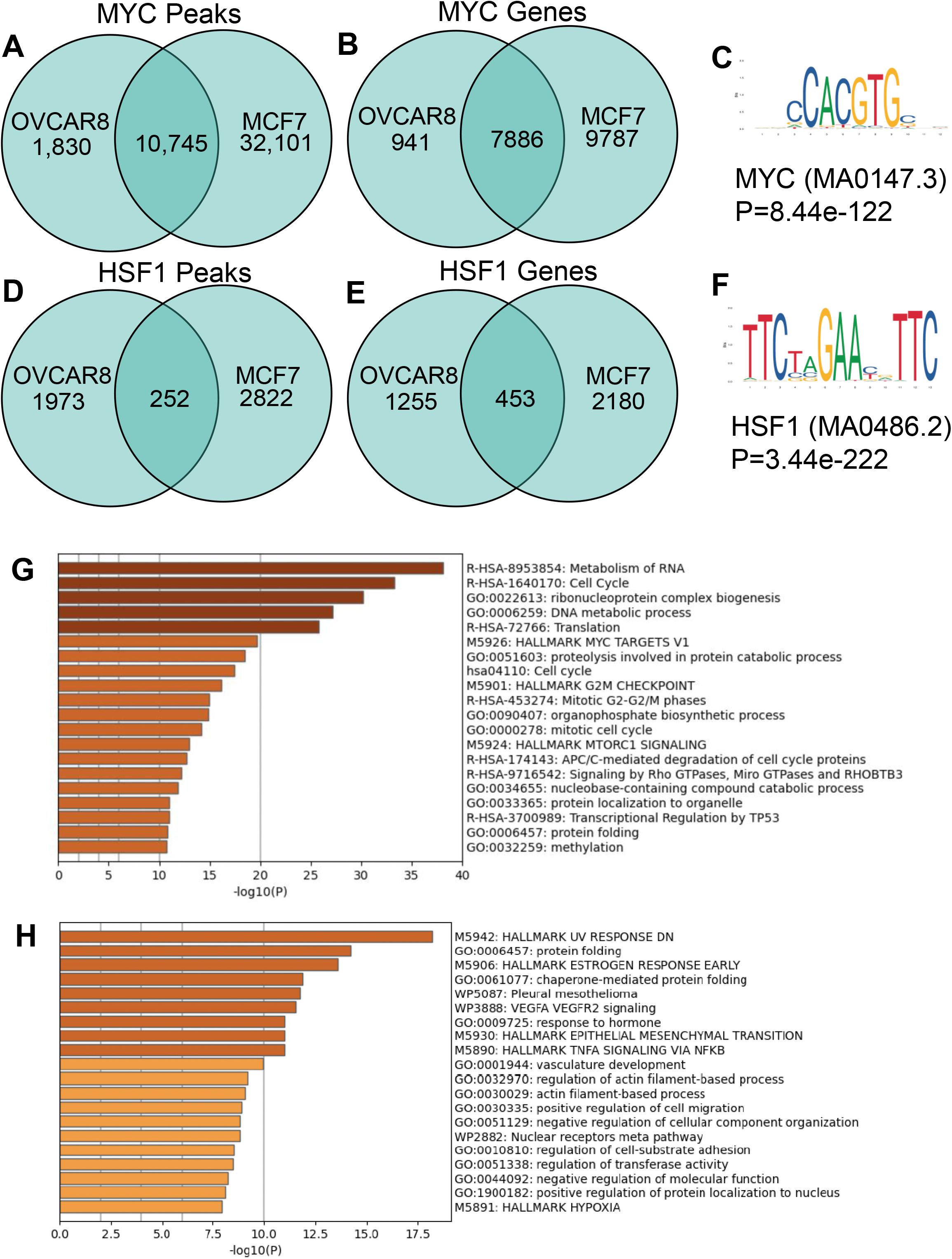
HSF1 and MYC CUT&RUN accurately detects their binding. A-B) MYC CUT&RUN peaks (A) and annotated genes (B) are compared to a previous MYC ChIP-Seq in MCF7 cells. C) The MYC motif that was detected from the MYC CUT&RUN. D-E) HSF1 CUT&RUN peaks (A) and annotated genes (B) are compared to a previous HSF1 ChIP-Seq in MCF7 cells. F) The HSF1 motif that was detected from the HSF1 CUT&RUN. G-H) Gene ontology for annotated genes for MYC (G) and HSF1 (H) binding.

**Supplemental Figure 4:**
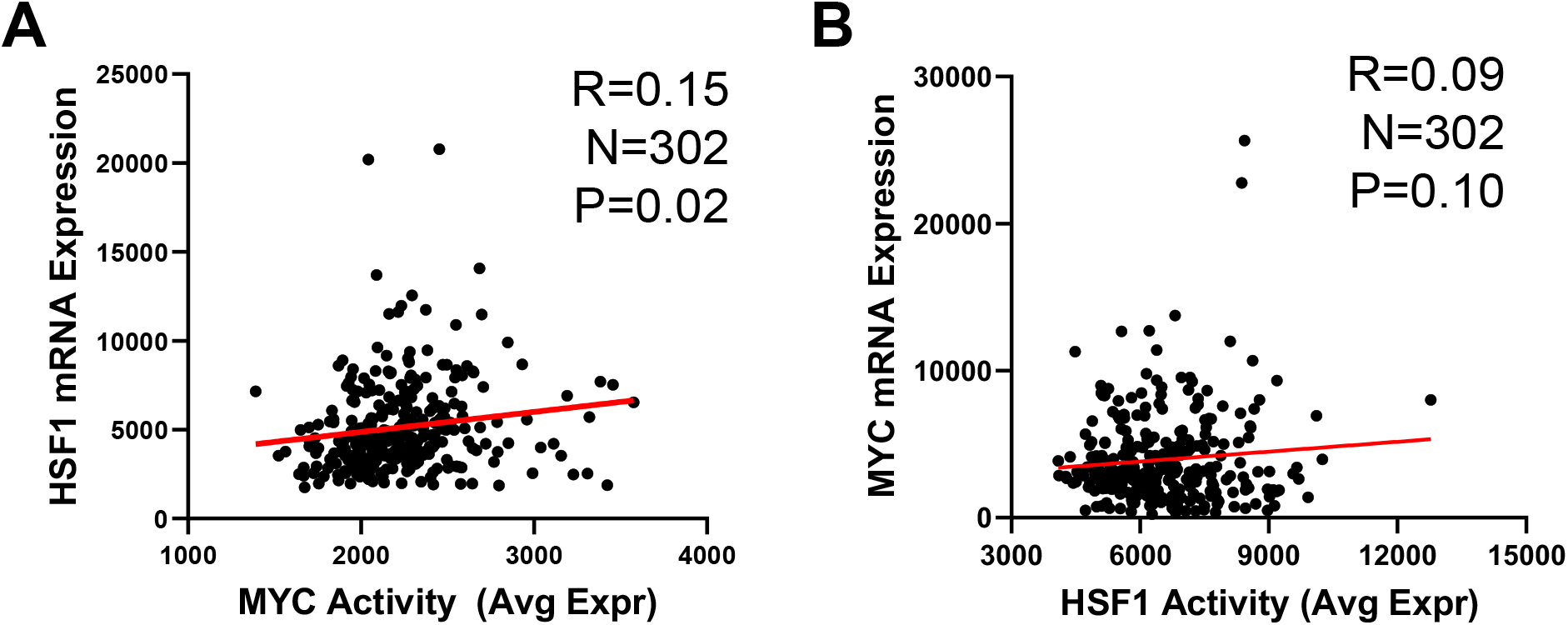
Correlation of MYC and HSF1 activity with their expression. A-B) MYC (A) and HSF1 (B) activity was assessed using published gene signatures and subjected to a Pearson correlation assessing the relationship of MYC activity with HSF1 expression (A) and HSF1 activity with MYC expression (B).

**Supplemental Figure 5:**
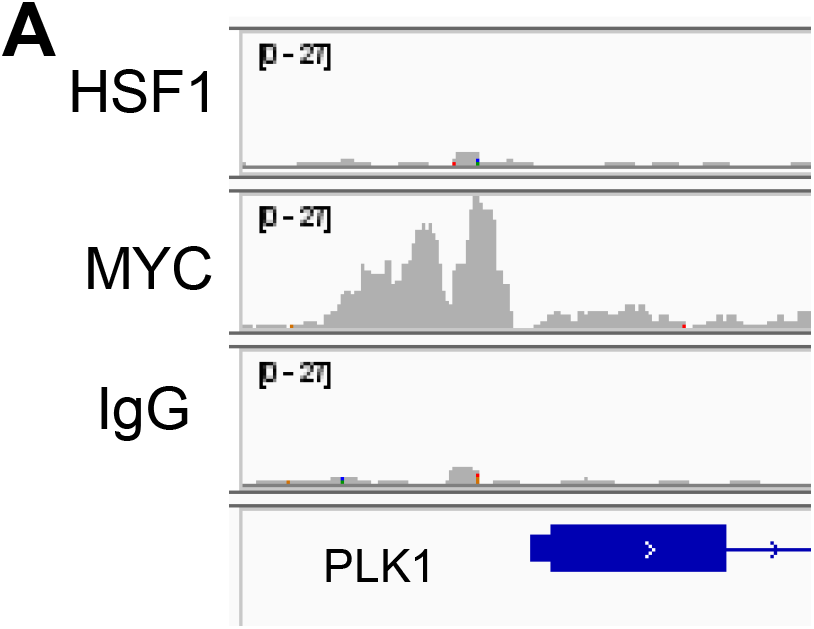
MYC binds the PLK1 gene promoter. Gene tracks HSF1 and MYC at the PLK1 promoter analyzed from CUT&RUN in Fig. 3.

## References

1. ACS. Cancer Facts & Figures 2024. Atlanta, GA2024.

2. Bowtell DD, Bohm S, Ahmed AA, Aspuria PJ, Bast RC, Jr., Beral V, et al. Rethinking ovarian cancer II: reducing mortality from high-grade serous ovarian cancer. Nat Rev Cancer 2015;15:668–79

3. Cress RD, Chen YS, Morris CR, Petersen M, Leiserowitz GS. Characteristics of Long-Term Survivors of Epithelial Ovarian Cancer. Obstet Gynecol 2015;126:491–7

4. Labidi-Galy SI, Papp E, Hallberg D, Niknafs N, Adleff V, Noe M, et al. High grade serous ovarian carcinomas originate in the fallopian tube. Nat Commun 2017;8:1093

5. Lheureux S, Gourley C, Vergote I, Oza AM. Epithelial ovarian cancer. Lancet 2019;393:1240–53

6. Torre LA, Trabert B, DeSantis CE, Miller KD, Samimi G, Runowicz CD, et al. Ovarian cancer statistics, 2018. CA Cancer J Clin 2018;68:284–96

7. Lengyel E. Ovarian cancer development and metastasis. The American journal of pathology 2010;177:1053–64

8. Dudler R, Travers AA. Upstream elements necessary for optimal function of the hsp 70 promoter in transformed flies. Cell 1984;38:391–8

9. Parker CS, Topol J. A Drosophila RNA polymerase II transcription factor contains a promoter-region-specific DNA-binding activity. Cell 1984;36:357–69

10. Topol J, Ruden DM, Parker CS. Sequences required for in vitro transcriptional activation of a Drosophila hsp 70 gene. Cell 1985;42:527–37

11. Gomez-Pastor R, Burchfiel ET, Thiele DJ. Regulation of heat shock transcription factors and their roles in physiology and disease. Nature reviews Molecular cell biology 2018;19:4–19

12. Jacobs C, Shah S, Lu WC, Ray H, Wang J, Hockaden N, et al. HSF1 Inhibits Antitumor Immune Activity in Breast Cancer by Suppressing CCL5 to Block CD8+ T-cell Recruitment. Cancer Res 2024;84:276–90

13. Cen H, Zheng S, Fang YM, Tang XP, Dong Q. Induction of HSF1 expression is associated with sporadic colorectal cancer. World journal of gastroenterology 2004;10:3122–6

14. Chen YF, Wang SY, Yang YH, Zheng J, Liu T, Wang L. Targeting HSF1 leads to an antitumor effect in human epithelial ovarian cancer. International journal of molecular medicine 2017;39:1564–70

15. Cheng Q, Chang JT, Geradts J, Neckers LM, Haystead T, Spector NL, et al. Amplification and high-level expression of heat shock protein 90 marks aggressive phenotypes of human epidermal growth factor receptor 2 negative breast cancer. Breast cancer research : BCR 2012;14:R62

16. Chuma M, Sakamoto N, Nakai A, Hige S, Nakanishi M, Natsuizaka M, et al. Heat shock factor 1 accelerates hepatocellular carcinoma development by activating nuclear factor-kappaB/mitogen-activated protein kinase. Carcinogenesis 2014;35:272–81

17. Cui J, Tian H, Chen G. Upregulation of Nuclear Heat Shock Factor 1 Contributes to Tumor Angiogenesis and Poor Survival in Patients With Non-Small Cell Lung Cancer. The Annals of thoracic surgery 2015;100:465–72

18. Dai C, Whitesell L, Rogers AB, Lindquist S. Heat shock factor 1 is a powerful multifaceted modifier of carcinogenesis. Cell 2007;130:1005–18

19. Dudeja V, Chugh RK, Sangwan V, Skube SJ, Mujumdar NR, Antonoff MB, et al. Prosurvival role of heat shock factor 1 in the pathogenesis of pancreatobiliary tumors. American journal of physiology Gastrointestinal and liver physiology 2011;300:G948–55

20. Engerud H, Tangen IL, Berg A, Kusonmano K, Halle MK, Oyan AM, et al. High level of HSF1 associates with aggressive endometrial carcinoma and suggests potential for HSP90 inhibitors. British journal of cancer 2014;111:78–84

21. Fang F, Chang R, Yang L. Heat shock factor 1 promotes invasion and metastasis of hepatocellular carcinoma in vitro and in vivo. Cancer 2012;118:1782–94

22. Hoang AT, Huang J, Rudra-Ganguly N, Zheng J, Powell WC, Rabindran SK, et al. A novel association between the human heat shock transcription factor 1 (HSF1) and prostate adenocarcinoma. The American journal of pathology 2000;156:857–64

23. Ishiwata J, Kasamatsu A, Sakuma K, Iyoda M, Yamatoji M, Usukura K, et al. State of heat shock factor 1 expression as a putative diagnostic marker for oral squamous cell carcinoma. International journal of oncology 2012;40:47–52

24. Jego G, Lanneau D, De Thonel A, Berthenet K, Hazoume A, Droin N, et al. Dual regulation of SPI1/PU.1 transcription factor by heat shock factor 1 (HSF1) during macrophage differentiation of monocytes. Leukemia 2014;28:1676–86

25. Jin X, Moskophidis D, Mivechi NF. Heat shock transcription factor 1 is a key determinant of HCC development by regulating hepatic steatosis and metabolic syndrome. Cell metabolism 2011;14:91–103

26. Kim SA, Kwon SM, Yoon JH, Ahn SG. The antitumor effect of PLK1 and HSF1 double knockdown on human oral carcinoma cells. International journal of oncology 2010;36:867–72

27. Kourtis N, Moubarak RS, Aranda-Orgilles B, Lui K, Aydin IT, Trimarchi T, et al. FBXW7 modulates cellular stress response and metastatic potential through HSF1 post-translational modification. Nature cell biology 2015;17:322–32

28. Li S, Ma W, Fei T, Lou Q, Zhang Y, Cui X, et al. Upregulation of heat shock factor 1 transcription activity is associated with hepatocellular carcinoma progression. Molecular medicine reports 2014;10:2313–21

29. Mendillo ML, Santagata S, Koeva M, Bell GW, Hu R, Tamimi RM, et al. HSF1 drives a transcriptional program distinct from heat shock to support highly malignant human cancers. Cell 2012;150:549–62

30. Min JN, Huang L, Zimonjic DB, Moskophidis D, Mivechi NF. Selective suppression of lymphomas by functional loss of Hsf1 in a p53-deficient mouse model for spontaneous tumors. Oncogene 2007;26:5086–97

31. Nakamura Y, Fujimoto M, Hayashida N, Takii R, Nakai A, Muto M. Silencing HSF1 by short hairpin RNA decreases cell proliferation and enhances sensitivity to hyperthermia in human melanoma cell lines. Journal of dermatological science 2010;60:187–92

32. Powell CD, Paullin TR, Aoisa C, Menzie CJ, Ubaldini A, Westerheide SD. The Heat Shock Transcription Factor HSF1 Induces Ovarian Cancer Epithelial-Mesenchymal Transition in a 3D Spheroid Growth Model. PloS one 2016;11:e0168389

33. Santagata S, Hu R, Lin NU, Mendillo ML, Collins LC, Hankinson SE, et al. High levels of nuclear heat-shock factor 1 (HSF1) are associated with poor prognosis in breast cancer. Proceedings of the National Academy of Sciences of the United States of America 2011;108:18378–83

34. Tsukao Y, Yamasaki M, Miyazaki Y, Makino T, Takahashi T, Kurokawa Y, et al. Overexpression of heat-shock factor 1 is associated with a poor prognosis in esophageal squamous cell carcinoma. Oncology letters 2017;13:1819–25

35. Wu PS, Chang YH, Pan CC. High expression of heat shock proteins and heat shock factor-1 distinguishes an aggressive subset of clear cell renal cell carcinoma. Histopathology 2017;71:711–8

36. Yasuda K, Hirohashi Y, Mariya T, Murai A, Tabuchi Y, Kuroda T, et al. Phosphorylation of HSF1 at serine 326 residue is related to the maintenance of gynecologic cancer stem cells through expression of HSP27. Oncotarget 2017;8:31540–53

37. Zhang N, Wu Y, Lyu X, Li B, Yan X, Xiong H, et al. HSF1 upregulates ATG4B expression and enhances epirubicin-induced protective autophagy in hepatocellular carcinoma cells. Cancer letters 2017;409:81–90

38. Zhou Z, Li Y, Jia Q, Wang Z, Wang X, Hu J, et al. Heat shock transcription factor 1 promotes the proliferation, migration and invasion of osteosarcoma cells. Cell proliferation 2017;50

39. Carpenter RL, Sirkisoon S, Zhu D, Rimkus T, Harrison A, Anderson A, et al. Combined inhibition of AKT and HSF1 suppresses breast cancer stem cells and tumor growth. Oncotarget 2017;8:73947–63

40. Gökmen-Polar Y, Badve S. Upregulation of HSF1 in estrogen receptor positive breast cancer. Oncotarget 2016;7:84239–45

41. Chen CH, Shen J, Lee WJ, Chow SN. Overexpression of cyclin D1 and c-Myc gene products in human primary epithelial ovarian cancer. Int J Gynecol Cancer 2005;15:878–83

42. Baker VV, Borst MP, Dixon D, Hatch KD, Shingleton HM, Miller D. c-myc amplification in ovarian cancer. Gynecol Oncol 1990;38:340–2

43. Buchynska LG, Brieieva OV, Iurchenko NP. Assessment of HER-2/neu, small es, Cyrillic-MYC and CCNE1 gene copy number variations and protein expression in endometrial carcinomas. Exp Oncol 2019;41:138–43

44. Jung M, Russell AJ, Kennedy C, Gifford AJ, Mallitt KA, Sivarajasingam S, et al. Clinical Importance of Myc Family Oncogene Aberrations in Epithelial Ovarian Cancer. JNCI Cancer Spectr 2018;2:pky047

45. Li C, Bonazzoli E, Bellone S, Choi J, Dong W, Menderes G, et al. Mutational landscape of primary, metastatic, and recurrent ovarian cancer reveals c-MYC gains as potential target for BET inhibitors. Proceedings of the National Academy of Sciences of the United States of America 2019;116:619–24

46. Papp E, Hallberg D, Konecny GE, Bruhm DC, Adleff V, Noe M, et al. Integrated Genomic, Epigenomic, and Expression Analyses of Ovarian Cancer Cell Lines. Cell Rep 2018;25:2617–33

47. Tian X, Song J, Zhang X, Yan M, Wang S, Wang Y, et al. MYC-regulated pseudogene HMGA1P6 promotes ovarian cancer malignancy via augmenting the oncogenic HMGA1/2. Cell Death Dis 2020;11:167

48. CGARN. Integrated genomic analyses of ovarian carcinoma. Nature 2011;474:609–15

49. Meyer N, Penn LZ. Reflecting on 25 years with MYC. Nat Rev Cancer 2008;8:976–90

50. Iba T, Kigawa J, Kanamori Y, Itamochi H, Oishi T, Simada M, et al. Expression of the c-myc gene as a predictor of chemotherapy response and a prognostic factor in patients with ovarian cancer. Cancer Sci 2004;95:418–23

51. Colombo I, Garg S, Danesh A, Bruce J, Shaw P, Tan Q, et al. Heterogeneous alteration of the ERBB3-MYC axis associated with MEK inhibitor resistance in a KRAS-mutated low-grade serous ovarian cancer patient. Cold Spring Harb Mol Case Stud 2019;5

52. Sheng Q, Zhang Y, Wang Z, Ding J, Song Y, Zhao W. Cisplatin-mediated down-regulation of miR-145 contributes to up-regulation of PD-L1 via the c-Myc transcription factor in cisplatin-resistant ovarian carcinoma cells. Clin Exp Immunol 2020;200:45–52

53. Yi J, Liu C, Tao Z, Wang M, Jia Y, Sang X, et al. MYC status as a determinant of synergistic response to Olaparib and Palbociclib in ovarian cancer. EBioMedicine 2019;43:225–37

54. Zeng M, Kwiatkowski NP, Zhang T, Nabet B, Xu M, Liang Y, et al. Targeting MYC dependency in ovarian cancer through inhibition of CDK7 and CDK12/13. Elife 2018;7

55. Xu M, Lin L, Ram BM, Shriwas O, Chuang KH, Dai S, et al. Heat shock factor 1 (HSF1) specifically potentiates c-MYC-mediated transcription independently of the canonical heat shock response. Cell Rep 2023;42:112557

56. Cigliano A, Pilo MG, Li L, Latte G, Szydlowska M, Simile MM, et al. Deregulated c-Myc requires a functional HSF1 for experimental and human hepatocarcinogenesis. Oncotarget 2017;8:90638–50

57. Hamanaka R, Maloid S, Smith MR, O’Connell CD, Longo DL, Ferris DK. Cloning and characterization of human and murine homologues of the Drosophila polo serine-threonine kinase. Cell Growth Differ 1994;5:249–57

58. Chiappa M, Petrella S, Damia G, Broggini M, Guffanti F, Ricci F. Present and Future Perspective on PLK1 Inhibition in Cancer Treatment. Front Oncol 2022;12:903016

59. Kressin M, Fietz D, Becker S, Strebhardt K. Modelling the Functions of Polo-Like Kinases in Mice and Their Applications as Cancer Targets with a Special Focus on Ovarian Cancer. Cells 2021;10

60. Zitouni S, Nabais C, Jana SC, Guerrero A, Bettencourt-Dias M. Polo-like kinases: structural variations lead to multiple functions. Nature reviews Molecular cell biology 2014;15:433–52

61. Liu Z, Sun Q, Wang X. PLK1, A Potential Target for Cancer Therapy. Transl Oncol 2017;10:22–32

62. Raab M, Sanhaji M, Zhou S, Rödel F, El-Balat A, Becker S, et al. Blocking Mitotic Exit of Ovarian Cancer Cells by Pharmaceutical Inhibition of the Anaphase-Promoting Complex Reduces Chromosomal Instability. Neoplasia 2019;21:363–75

63. Rödel F, Zhou S, Győrffy B, Raab M, Sanhaji M, Mandal R, et al. The Prognostic Relevance of the Proliferation Markers Ki-67 and Plk1 in Early-Stage Ovarian Cancer Patients With Serous, Low-Grade Carcinoma Based on mRNA and Protein Expression. Front Oncol 2020;10:558932

64. Takai N, Yoshimatsu J, Nishida Y, Narahara H, Miyakawa I, Hamanaka R. Expression of polo-like kinase (PLK) in the mouse placenta and ovary. Reprod Fertil Dev 1999;11:31–5

65. Weichert W, Denkert C, Schmidt M, Gekeler V, Wolf G, Köbel M, et al. Polo-like kinase isoform expression is a prognostic factor in ovarian carcinoma. British journal of cancer 2004;90:815–21

66. Zhang H, Zhang K, Xu Z, Chen Z, Wang Q, Wang C, et al. MicroRNA-545 suppresses progression of ovarian cancer through mediating PLK1 expression by a direct binding and an indirect regulation involving KDM4B-mediated demethylation. BMC Cancer 2021;21:163

67. Zhang R, Shi H, Ren F, Liu H, Zhang M, Deng Y, et al. Misregulation of polo-like protein kinase 1, P53 and P21WAF1 in epithelial ovarian cancer suggests poor prognosis. Oncol Rep 2015;33:1235–42

68. Zhang S, Jing Y, Zhang M, Zhang Z, Ma P, Peng H, et al. Stroma-associated master regulators of molecular subtypes predict patient prognosis in ovarian cancer. Sci Rep 2015;5:16066

69. Choi BH, Pagano M, Dai W. Plk1 protein phosphorylates phosphatase and tensin homolog (PTEN) and regulates its mitotic activity during the cell cycle. J Biol Chem 2014;289:14066–74

70. Lee YJ, Kim EH, Lee JS, Jeoung D, Bae S, Kwon SH, et al. HSF1 as a mitotic regulator: phosphorylation of HSF1 by Plk1 is essential for mitotic progression. Cancer Res 2008;68:7550–60

71. Li Z, Li J, Bi P, Lu Y, Burcham G, Elzey BD, et al. Plk1 phosphorylation of PTEN causes a tumor-promoting metabolic state. Mol Cell Biol 2014;34:3642–61

72. Liu XS, Song B, Elzey BD, Ratliff TL, Konieczny SF, Cheng L, et al. Polo-like kinase 1 facilitates loss of Pten tumor suppressor-induced prostate cancer formation. J Biol Chem 2011;286:35795–800

73. Ren Y, Bi C, Zhao X, Lwin T, Wang C, Yuan J, et al. PLK1 stabilizes a MYC-dependent kinase network in aggressive B cell lymphomas. J Clin Invest 2018;128:5517–30

74. Gatza ML, Lucas JE, Barry WT, Kim JW, Wang Q, Crawford MD, et al. A pathway-based classification of human breast cancer. Proceedings of the National Academy of Sciences of the United States of America 2010;107:6994–9

75. Karst AM, Levanon K, Drapkin R. Modeling high-grade serous ovarian carcinogenesis from the fallopian tube. Proceedings of the National Academy of Sciences of the United States of America 2011;108:7547–52

76. Blackman A, Rees AC, Bowers RR, Jones CM, Vaena SG, Clark MA, et al. MYC is sufficient to generate mid-life high-grade serous ovarian and uterine serous carcinomas in a p53-R270H mouse model. bioRxiv 2024:2024.01.24.576924

77. Lu WC, Omari R, Ray H, Wang J, Williams I, Jacobs C, et al. AKT1 mediates multiple phosphorylation events that functionally promote HSF1 activation. Febs j 2022;289:3876–93

78. Langmead B, Salzberg SL. Fast gapped-read alignment with Bowtie 2. Nature Methods 2012;9:357–9

79. Feng J, Liu T, Qin B, Zhang Y, Liu XS. Identifying ChIP-seq enrichment using MACS. Nature Protocols 2012;7:1728–40

80. McLeay RC, Bailey TL. Motif Enrichment Analysis: a unified framework and an evaluation on ChIP data. BMC Bioinformatics 2010;11:165

81. Hahne F, Ivanek R. Visualizing Genomic Data Using Gviz and Bioconductor. In: Mathé E, Davis S, editors. Statistical Genomics: Methods and Protocols. New York, NY: Springer New York; 2016. p 335–51.

82. Zhou Y, Zhou B, Pache L, Chang M, Khodabakhshi AH, Tanaseichuk O, et al. Metascape provides a biologist-oriented resource for the analysis of systems-level datasets. Nat Commun 2019;10:1523

83. Bankhead P, Loughrey MB, Fernández JA, Dombrowski Y, McArt DG, Dunne PD, et al. QuPath: Open source software for digital pathology image analysis. Sci Rep 2017;7:16878

84. Wang W, Fang F, Ozes A, Nephew KP. Targeting Ovarian Cancer Stem Cells by Dual Inhibition of HOTAIR and DNA Methylation. Mol Cancer Ther 2021

85. Brusselaers N, Ekwall K, Durand-Dubief M. Copy number of 8q24.3 drives HSF1 expression and patient outcome in cancer: an individual patient data meta-analysis. Hum Genomics 2019;13:54

86. Zhang CQ, Williams H, Prince TL, Ho ES. Overexpressed HSF1 cancer signature genes cluster in human chromosome 8q. Hum Genomics 2017;11:35

87. Chandriani S, Frengen E, Cowling VH, Pendergrass SA, Perou CM, Whitfield ML, et al. A core MYC gene expression signature is prominent in basal-like breast cancer but only partially overlaps the core serum response. PloS one 2009;4:e6693

88. Lee BK, Bhinge AA, Battenhouse A, McDaniell RM, Liu Z, Song L, et al. Cell-type specific and combinatorial usage of diverse transcription factors revealed by genome-wide binding studies in multiple human cells. Genome Res 2012;22:9–24

89. Dong B, Jaeger AM, Hughes PF, Loiselle DR, Hauck JS, Fu Y, et al. Targeting therapy-resistant prostate cancer via a direct inhibitor of the human heat shock transcription factor 1. Sci Transl Med 2020;12

90. Wang D, Pierce A, Veo B, Fosmire S, Danis E, Donson A, et al. A Regulatory Loop of FBXW7-MYC-PLK1 Controls Tumorigenesis of MYC-Driven Medulloblastoma. Cancers (Basel) 2021;13

91. Schaefer CF, Anthony K, Krupa S, Buchoff J, Day M, Hannay T, et al. PID: the Pathway Interaction Database. Nucleic Acids Res 2009;37:D674–9

92. Rudolph D, Steegmaier M, Hoffmann M, Grauert M, Baum A, Quant J, et al. BI 6727, a Polo-like kinase inhibitor with improved pharmacokinetic profile and broad antitumor activity. Clin Cancer Res 2009;15:3094–102

93. Pujade-Lauraine E, Selle F, Weber B, Ray-Coquard IL, Vergote I, Sufliarsky J, et al. Volasertib Versus Chemotherapy in Platinum-Resistant or -Refractory Ovarian Cancer: A Randomized Phase II Groupe des Investigateurs Nationaux pour l’Etude des Cancers de l’Ovaire Study. J Clin Oncol 2016;34:706–13

94. Xie FF, Pan SS, Ou RY, Zheng ZZ, Huang XX, Jian MT, et al. Volasertib suppresses tumor growth and potentiates the activity of cisplatin in cervical cancer. Am J Cancer Res 2015;5:3548–59

95. Spänkuch B, Matthess Y, Knecht R, Zimmer B, Kaufmann M, Strebhardt K. Cancer inhibition in nude mice after systemic application of U6 promoter-driven short hairpin RNAs against PLK1. J Natl Cancer Inst 2004;96:862–72

96. Spänkuch-Schmitt B, Bereiter-Hahn J, Kaufmann M, Strebhardt K. Effect of RNA silencing of polo-like kinase-1 (PLK1) on apoptosis and spindle formation in human cancer cells. J Natl Cancer Inst 2002;94:1863–77

97. Spänkuch-Schmitt B, Wolf G, Solbach C, Loibl S, Knecht R, Stegmüller M, et al. Downregulation of human polo-like kinase activity by antisense oligonucleotides induces growth inhibition in cancer cells. Oncogene 2002;21:3162–71

98. Kobayashi Y, Yamauchi T, Kiyoi H, Sakura T, Hata T, Ando K, et al. Phase I trial of volasertib, a Polo-like kinase inhibitor, in Japanese patients with acute myeloid leukemia. Cancer Sci 2015;106:1590–5

99. Ottmann OG, Müller-Tidow C, Krämer A, Schlenk RF, Lübbert M, Bug G, et al. Phase I dose-escalation trial investigating volasertib as monotherapy or in combination with cytarabine in patients with relapsed/refractory acute myeloid leukaemia. Br J Haematol 2019;184:1018–21

100. Ellis PM, Leighl NB, Hirsh V, Reaume MN, Blais N, Wierzbicki R, et al. A Randomized, Open-Label Phase II Trial of Volasertib as Monotherapy and in Combination With Standard-Dose Pemetrexed Compared With Pemetrexed Monotherapy in Second-Line Treatment for Non-Small-Cell Lung Cancer. Clin Lung Cancer 2015;16:457–65

101. Stadler WM, Vaughn DJ, Sonpavde G, Vogelzang NJ, Tagawa ST, Petrylak DP, et al. An open-label, single-arm, phase 2 trial of the Polo-like kinase inhibitor volasertib (BI 6727) in patients with locally advanced or metastatic urothelial cancer. Cancer 2014;120:976–82

102. Zhu W, Niu J, He M, Zhang L, Lv X, Liu F, et al. SNORD89 promotes stemness phenotype of ovarian cancer cells by regulating Notch1-c-Myc pathway. J Transl Med 2019;17:259

103. Lancho O, Herranz D. The MYC Enhancer-ome: Long-Range Transcriptional Regulation of MYC in Cancer. Trends Cancer 2018;4:810–22

104. Shen Q, Wang R, Liu X, Song P, Zheng M, Ren X, et al. HSF1 Stimulates Glutamine Transport by Super-Enhancer-Driven lncRNA LINC00857 in Colorectal Cancer. Cancers (Basel) 2022;14

105. Astrinidis A, Senapedis W, Henske EP. Hamartin, the tuberous sclerosis complex 1 gene product, interacts with polo-like kinase 1 in a phosphorylation-dependent manner. Hum Mol Genet 2006;15:287–97

106. Li Y, Wang H, Zhang Z, Tang C, Zhou X, Mohan C, et al. Identification of polo-like kinase 1 as a therapeutic target in murine lupus. Clin Transl Immunology 2022;11:e1362

107. Carpenter RL, Paw I, Dewhirst MW, Lo HW. Akt phosphorylates and activates HSF-1 independent of heat shock, leading to Slug overexpression and epithelial-mesenchymal transition (EMT) of HER2-overexpressing breast cancer cells. Oncogene 2015;34:546–57

108. Sodi VL, Khaku S, Krutilina R, Schwab LP, Vocadlo DJ, Seagroves TN, et al. mTOR/MYC Axis Regulates O-GlcNAc Transferase Expression and O-GlcNAcylation in Breast Cancer. Mol Cancer Res 2015;13:923–33

109. Yeh CH, Bellon M, Nicot C. FBXW7: a critical tumor suppressor of human cancers. Mol Cancer 2018;17:115

110. Zhang T, Li N, Sun C, Jin Y, Sheng X. MYC and the unfolded protein response in cancer: synthetic lethal partners in crime? EMBO Mol Med 2020;12:e11845

111. Pasqua AE, Sharp SY, Chessum NEA, Hayes A, Pellegrino L, Tucker MJ, et al. HSF1 Pathway Inhibitor Clinical Candidate (CCT361814/NXP800) Developed from a Phenotypic Screen as a Potential Treatment for Refractory Ovarian Cancer and Other Malignancies. J Med Chem 2023;66:5907–36

